# Medium optimization for biomass production of three peat moss (*Sphagnum* L.) species using fractional factorial design and response surface methodology

**DOI:** 10.1101/2021.03.19.436135

**Authors:** Melanie A. Heck, Ingrida Melková, Clemens Posten, Eva L. Decker, Ralf Reski

**Affiliations:** Plant Biotechnology, Faculty of Biology, University of Freiburg, Freiburg, Germany; Institute of Process Engineering in Life Sciences III Bioprocess Engineering, Karlsruhe Institute of Technology (KIT), Karlsruhe, Germany; CIBSS – Centre for Integrative Biological Signalling Studies, University of Freiburg, Freiburg, Germany; Cluster of Excellence livMatS @ FIT – Freiburg Center for Interactive Materials and Bioinspired Technologies, University of Freiburg, Freiburg, Germany

**Keywords:** Design of Experiments, medium optimization, peat moss, photobioreactor, Sphagnum farming

## Abstract

Peat moss (*Sphagnum*) biomass is a promising bioresource to substitute peat in growing media with a renewable material. For sustainable production on a large scale, the productivity of *Sphagnum* mosses has to be increased by optimizing culture conditions. Optimization was achieved using fractional factorial design and response surface methodology based on central composite design to determine concentrations of eight factors leading to highest biomass yield. We improved a standard Sphagnum medium by reducing the concentrations of NH_4_NO_3_, KH_2_PO_4_, KCl, MgSO_4_, Ca(NO_3_)_2_, FeSO_4_ and a microelement solution up to 50 %. Together with a reduced sucrose concentration for *Sphagnum fuscum*, while it remained unchanged for *Sphagnum palustre* and *Sphagnum squarrosum*, moss productivities were enhanced for all tested species in shake flasks. Further upscaling to 5 L photobioreactors increased the biomass yield up to nearly 50-fold for *S. fuscum*, 40-fold for *S. palustre* and 25-fold for *S. squarrosum* in 24 days.

## 1. Introduction

Peat mosses (*Sphagnum* spec.) are among the oldest land plants and are rapidly gaining interest in basic and applied research. Their aplications range from traditional medicine like wound dressing (Sabovljević et al., 2016) to biotechnological applications like biomonitoring of air pollutants (Aboal et al., 2020; Capozzi et al., 2017; Di Palma et al., 2019). In addition, they are an important climate change factor as a major constituent of peatlands, which are the largest terrestrial long-term carbon storage (Joosten et al., 2016). Peatlands cover around 3 % of the world land area, but store around a quarter of the world’s soil carbon (Turetsky et al., 2015). Therefore, peat mosses play an important role in earth’s climate and have a large impact on global carbon cycling, which makes them a suitable plant model in carbon cycling studies (Weston et al., 2018). The availability of the first genome sequences of two *Sphagnum* species (*Sphagnum fallax* v1.1 and *Sphagnum magellanicum* v1.1, DOE-JGI, http://phytozome-next.jgi.doe.gov/) will further extend the scientific impact and research possibilities, as has been observed for the model moss *Physcomitrella patens* (Rensing et al., 2008).

Over the long term, the peatland subsistence base is getting destroyed for peat extraction or their agricultural use. Drainage leads to release of the stored carbon, which accounts for 32 % of global cropland greenhouse gas (GHG) emission (Carlson et al., 2017). However, *Sphagnum* biomass can be produced in an environmentally friendly and sustainable land use option on rewetted peatlands, an application named Sphagnum farming (Gaudig et al., 2014). Rewetting drained peatlands reduces GHG emisions and simultaneously produces a renewable alternative to substitute fossil peat, which is the best-quality horticultural growing medium so far (Gaudig et al., 2018). The suitability for their use in these growing media have tested positively for various species, e.g. *S. fuscum* (section *Acutifolia*), *S. palustre* (section *Sphagnum*) or *S. squarrosum* (section *Squarrosa*), depending on the plant under cultivation and proportion of the biomass in the culture substrate (Gaudig et al., 2018).

The large-scale implementation of Sphagnum farming is limited by the lack of peat moss as a founder material. So far, there is no supply for *Sphagnum* due to the scarcity and conservation status of *Sphagnum* mosses, for example by the European Council Habitats Directive (92/43/EEC). In addition, peat moss collected in natural habitats may include undesired *Sphagnum* species or vascular plants which could worsen or limit its use (Gaudig et al., 2018). A promising method for providing the required founder material is based on vegetative regeneration of *Sphagnum* under controlled conditions, since peat mosses regenerate from many parts of the shoot like capitula, branches and stems, but not from leaves (Poschlod and Pfadenhauer, 1989; Sobotka, 2015). While *Sphagnum* propagates slowly under natural conditions, i*n-vitro* cultivation could accelerate the peat moss growth.

An important step towards the large-scale production of such founder material was the development of an axenic photobioreactor production process on a laboratory scale, e.g. for *Sphagnum palustre* with a 30-fold biomass increase within four weeks (Beike *et al*., 2015). This accelerated the production of founder material drastically compared to the 2.5-fold biomass increase per year in the field and in glasshouse experiments (Gaudig *et al*., 2014; Beike *et al*., 2015). Another important step was the establishment of axenic *in-vitro* cultures of 19 *Sphagnum* species with a selection of productive clones to achieve maximum yields (Heck et al., 2021), which was based on a medium developed for *S. palustre* by Beike *et al*. (2015). However, Heck *et al*. (2021) noted that this medium was not optimal for all of the tested species, because, for example, the productivity of *S. subnitens* as one of the most productive species from *Sphagnum*-dominated wetlands (Gunnarsson, 2005) had only a weak performance *in vitro*. Consequently, although the basic cultivation techniques are established, the cultivation process requires optimization for each *Sphagnum* species.

In the context of medium optimization, Design of Experiments (DOE) is a powerful statistical tool for product and process design, development and optimization. DOE gained an increasing interest in all scientific areas over traditional methods like one factor at time (OFAT) (Duraković, 2017). Changing one factor, while keeping the others constant, is a time-consuming process, as it requires a high number of experiments without determining the existence of interactions between individual factors. DOE overcomes these limitations by providing better results with fewer experiments (Fukuda et al., 2018). DOE helps to determine the most important input factors, understand the interaction between factors and identify the factor settings leading to optimized output responses (Fukuda et al., 2018). Experimental designs can be divided into two steps: 1) screening designs, like fractional factorial design at two levels, where many factors are studied to identify the significant ones and 2) optimization designs, like the central composite as one of the most widely used designs, where the factors are further examined to determine the best conditions (Candioti et al., 2014; Singh et al., 2017; Maina et al., 2019).

Here, we report on the optimized media compositions for peat moss biomass production, obtained by the screening of eight factors using two-level fractional factorial design and optimization of these factors by central composite design for axenic *in-vitro* cultivation of the three species *S. fuscum*, *S. palustre* and *S. squarrosum* in small scale. These species are favourable bioresource candidates for both Sphagnum farming and growing media, with *S. palustre* one of the most promising peat mosses (Gaudig et al., 2014). The results from an optimized 5 L laboratory scale photobioreactor process for all three species is presented also, which may serve as basis for a large scale process for peat moss biomass production.

## 2. Materials and methods

### 2.1. *In-vitro* cultivation

The peat moss strains for this study each derived from a single spore, are described in Heck et al. (2021), and are vailable from the International Moss Stock Center (https://www.moss-stock-center.org) under their respective IMSC accession number. For suspension cultures of the clones *S. fuscum* 1.1 (IMSC #41158), *S. palustre* 12a (IMSC #40068) and *S. squarrosum* 5.2 (IMSC #41193), gametophores were disrupted with forceps in laminar flow benches (LaminAir, Heraeus, Hanau, Germany) and transferred to 500 or 1000 mL Erlenmeyer flasks filled with 200 or 500 mL liquid Sphagnum medium, respectively. Standard Sphagnum medium consists of Knop medium (1.84 mM KH_2_PO_4_, 3.35 mM KCl, 1.01 mM MgSO_4_ · 7 H_2_O, 4.24 mM Ca(NO_3_)_2_ ·4 H_2_O, 45 µM FeSO_4_ · 7 H_2_O) according to Reski & Abel (1985), supplemented with microelements (ME) (50 µM H3BO3, 50 µM MnSO4 · H_2_O, 15 µM ZnSO4 · 7 H_2_O, 2.5 µM KJ, 500 nM Na2MoO4 · 2 H_2_O, 50 nM CuSO4 · 5 H_2_O, 50 nM Co(NO3)2 · 6 H_2_O) according to Schween *et al*. (2003), and supplemented with 2 % sucrose and 1.25 mM NH4NO3 with an adjusted pH of 4.8 before autoclaving according to Beike et al. (2015). The flasks were closed with Silicosen® silicone sponge plugs (Hirschmann Laborgeräte GmbH & Co. KG, Eberstadt, Germany) to allow gas exchange, and cultivated on a rotary shaker at 120 rpm (B. Braun Biotech International GmbH, Melsungen, Germany). Standard cultivation conditions were: climate chamber at a temperature of 22 °C under a photoperiod regime of 16/8 h (light/dark) with a light intensity of 70 ± 5 μmol m^−^²s^−1^ provided from above by fluorescent tubes (Master TL-D Super 80, Philips, Amsterdam, The Netherlands). Average measurements of light intensity were done manually with a planar quantum sensor (Li-Cor 250, Li-Cor Biosciences GmbH, Bad Homburg, Germany).

To check for putative contaminations, either medium or moss material were transferred to plates containing three different solid media, Knop ME, LB, and TSA, respectively. For detection of contamination, Knop ME medium was supplemented with 1 % glucose and 12 g L^−1^ Purified Agar (Oxoid Ltd. UK) with an adjusted pH of 5.8 before autoclaving. LB medium contained 10 g L^−1^ Bacto Tryptone (Becton, Dickinson & Co., NJ, USA), 10 g L^−1^ NaCl, 5 g L^−1^ Bacto Yeast Extract (Becton, Dickinson & Co.) and 15 g L^−1^ Bacto Agar (Becton, Dickinson & Co.) with an adjusted pH of 7.0 before autoclaving. Tryptic Soy Agar (TSA) contained 15 g L^−1^ peptone from casein, 5 g L^−1^ soy peptone, 5 g L^−1^ NaCl, 1 % glucose and 12 g L^−1^ purified agar (Oxoid Limited) with an adjusted pH of 7.5 before autoclaving. These control plates were sealed with Parafilm and incubated for four weeks at room temperature. A culture was considered axenic if no contamination on the plates occurred within that time.

### 2.2. Bioreactor cultivation

For scaling-up the cultivation process of *S. fuscum*, *S. palustre* and *S. squarrosum,* glass tank photobioreactors with 5.4 L working volume were used (Applikon Biotechnology, Schiedam, The Netherlands). A bioreactor was inoculated with a two-week-old preculture grown in Erlenmeyer flasks. For inoculum, 15 g of peat moss was weighed after filtering for 1 min using a Steritop filter and a vacuum pump, disrupted with forceps and filled in a flask containing 500 mL Sphagnum medium and cultivated under standard cultivation conditions. The whole content of the flask was transferred with 5 L of optimized medium to the photobioreactor. The reactor was iluminated with 12 LED stripes (MaxLine70, Lumitronix, Hechingen, Germany) placed around the reactor at 2 cm distance. The light intensity, measured behind the reactor glas wall, was increased stepwise from initially 150 µmol m^−2^ s^−1^ to 300 µmol m^−2^ s^−1^ at day 3 up to 500 µmol m^−2^ s^−1^ at day 7 with a day/night cycle of 20/4 h and aeration of 0.3 vvm with 2 % CO_2_-enriched air, which was passed trough a water bottle before entering the bioreactor. The bioreactor was equipped with a marine impeller placed 22 cm above the bottom. The cultures of *S. palustre* and *S. squarrosum* were stirred constantly with 100 rpm starting at day 17 to ensure that the moss plants did not aggregate. During the cultivation of *S. fuscum*, the bioreactor was not stirred, as no clumping of the mosses occurred. The pH was not adjusted, but was tracked during cultivation with an internal pH electrode (Applikon Biotechnology). The biomass increase was documented photographically directly after inoculation and at 3, 7, 10, 13, 15, 17, 20, 22 and 24 days. At the same time points 40 mL of medium were withdrawn from the bioreactor to measure nutrient consumption (see 2.3). After 24 days all material of the bioreactor was harvested and the biomass, fresh weights (see 2.4), and dry weights (see 2.5) were determined.

The absolute growth rate µ was calculated by the difference of the inoculum and the harvested fresh biomass: µ=(M_Harvest_-M_Inocula_) (t_Harvest_-t_Inocula_)^−1^, where M_Harvest_ is the harvested fresh biomass in mg FW L^−1^, M_Inocula_ is the inoculated biomass in mg FW L^−1^ and t the corresponding cultivation days.

### 2.3. Nutrient measurement

Prior to nutrient measurement all samples were passed through 0.45 µm PVDF filters (Rotalibo, Carl Roth, Karlsruhe, Germany). Inorganic ions were determined by an ion chromatograph (822 Compact IC plus, Methrom, Herisau, Switzerland) equipped with a Metrosep A Supp 5 150/4 column (Metrohm) and a guard column A Supp 4/5 Guard 4.0 (Metrohm) to determine anion concentration (Cl^−^, NO_3_, PO_4_, SO_4_). The eluent for the anion measurement was an aqueous solution of 3.2 mM Na_2_CO_3_, 1.0 mM NaHCO_3_ and 12.5 % (v/v) acetonitrile. To determine cation concentration (Na^+^, NH_4_^+^, K^+^, Ca^2+^, Mg^2+^), Metrosep C4 150/4.0 columns (Metrohm) and Metrosep C4 S-Gurad columns (Metrohm) with an eluent of 1.7 mM HNO_3_ and 0.7 mM 2,6-pyridinedicarboxylic acid were used. All solutions were prepared with ultra pure water (resistance 16 MΩ; Maxima, ELGA LabWater, Celle, Germany). Filtrated medium samples were diluted and injected automatically by an autosample unit (885 Professional sample Processor, Methrom) and analysed with a conductivity detector (Metrohm). This system was controlled and data processed using MagIC Net 2.3 software (Metrohm).

The analysis of glucose, fructose and sucrose was carried out with the Sucrose/D-Fructose/D-Glucose Assay Kit (Megazyme, Bray, Ireland) according to the manufacturer’s protocol. For use in 96 well microplates (Greiner Bio One, Kremsmünster, Austria) the assay volumes were reduced to 10 %. A sucrose and D-glucose/D-fructose standard curve was performed on each microplate and the results were calculated from the calibration curve. The absorbance was measured at 340 nm with a microplate reader (CLARIOstar, BMG Labtech, Ortenberg, Germany).

### 2.4. Fresh weight determination

Fresh weight was measured by filtering the total content of the bioreactor with a Büchner funnel and generating a vacuum for 1 minute by closing the funnel with a plug sealed with Parafilm. The amount of the peat moss biomass retained on the filter was weighed on a scale (E12000 S, Sartorius, Göttingen, Germany).

### 2.5. Dry weight determination

To measure dry weights from flask cultures, total biomass was harvested by filtering with a Büchner funnel and a vacuum pump. The moss material was transferred to pre-dried (0.5 h at 105 °C) aluminum weighing pans (Köhler Technische Produkte, Neulußheim, Germany) and dried for 2 hours at 105 °C in a forced air oven (Ehret GmbH Life Science Solutions, Freiburg, Germany). The dried moss material in the weighing pan was weighed with an accuracy scale (CPA 3245, Sartorius).

The moss material from the bioreactor (after fresh weight determination) was filled in a miracloth bag, dried for 10 h at 80 °C and weighed on a scale.

### 2.6. Design of experiments (DOE) for medium optimization

The optimization of the medium components for biomass increase was carried out for *S. fuscum*, *S. palustre* and *S. squarrosum*. The influence and significance of the media components, including the eight factors sucrose, NH_4_NO_3_, KH_2_PO_4_, KCl, MgSO_4_, Ca(NO_3_)_2_, ME and FeSO_4_, on the biomass yield (produced dry weight) were determined and optimized using Design-Expert® software (Version 11.1.2.0, Stat-Ease, Minneapolis, MN, USA) based on the variance analysis (ANOVA).

#### 2.6.1. Identification of important components

Screening was done using a two-level factorial design (2^k-p^), where low versus high factor settings were compared resulting in a linear model. To detect non-linearity, center points have to be added, located at the exact mid-point of all factor settings. Eight factors (k=8), which represent the main components of the standard Sphagnum medium, were selected. Each of the factors was set at three levels: low (−1), high (+1) and the center point inbetween (0). Sucrose range from 3 to 20 g L^−1^ and the other factors between 50 and 100 % of the standard Sphagnum medium for *S. fuscum* and *S. squarrosum* and between 10 and 100 % for *S. palustre* (Table 1). In the fractional factorial design, k factors were screened based on just 2^k-4^ experiments resulting in 16 runs conducted as duplicates and four replicates of the center point. In total 36 experiments were performed in random order. The flasks were filled with 200 mL of the respective medium and inoculated with 250 mg of moss material each. Before weighing the disrupted gametophores with an accuracy scale (E12000 S, Sartorius) in laminar flow benches, the gametophores were filtered for 1 min using a Steritop filter (Millipore Corporation, Billerica, MA, USA) and a vacuum pump (Vacuubrand MZ 2C, Vacuubrand GmbH and Co, Wertheim, Germany). After cultivation for four weeks under standard cultivation conditions, dry weights were determined and the significant factors identified.

**Table 1:** The factors and their levels for two-level factorial design as first screening of the optimized media composition towards biomass production of *S. fuscum*, *S. squarrosum* and *S. palustre*. Code level represents a change in concentration of each factor: −1 reduced concentration, +1 elevated concentration, 0 inbetween (center point).

| Symbols | Factor | Code Level |  |  |  |  |  |
| --- | --- | --- | --- | --- | --- | --- | --- |
|  |  | -1 | 0 | +1 | -1 | 0 | +1 |
|  |  | *S. fuscum/S. squarrosus* |  |  | *S. palustre* |  |  |
| A | Sucrose (g L<sup>-1</sup>) | 3 | 11.5 | 20 | 3 | 11.5 | 20 |
| B | NH<sub>4</sub>NO<sub>3</sub> (mM) | 0.63 | 0.94 | 1.25 | 0.5 | 0.875 | 1.25 |
| C | KH<sub>2</sub>PO<sub>4</sub> (mM) | 0.92 | 1.38 | 1.84 | 0.18 | 1.01 | 1.84 |
| D | KCl (mM) | 1.68 | 2.52 | 3.35 | 0.34 | 1.845 | 3.35 |
| E | MgSO<sub>4</sub> (mM) | 0.51 | 0.76 | 1.01 | 0.1 | 0.555 | 1.01 |
| F | Ca(NO<sub>3</sub>)<sub>2</sub> (mM) | 2.12 | 3.18 | 4.24 | 0.42 | 2.33 | 4.24 |
| G | ME (%) | 50 | 75 | 100 | 10 | 55 | 100 |
| H | FeSO<sub>4</sub> (μM) | 22.5 | 33.8 | 45 | 4.5 | 24.75 | 45 |

The results were analyzed by selecting the factors with the highest t-values. Effects below the t-value limit were only selected to support hierarchy. This means that a factor was selected even if it has a non-significant effect on the response, but interaction with another factor significantly influences response. In this design, all interaction effects are aliased, also known as confounded, as the number of experiments in the fractional factorial design is smaller than the number of different treatment combinations. For example, the effects of AB, CE, DH and FG are aliased and the effect of one of these combinations cannot be distinguished from the others.

The selected factorial model was checked using ANOVA. To trust the model, the following terms were checked: the p-value of the model term ≤ 0.05 reveals that the model is significant; the p-values of the selected factors ≤ 0.05 indicates that the factors are significant and have an effect on the response or they have to be selected as they are important for the interaction effects together with another factor. The lack of fit p-value > 0.05 indicates that the lack of fit is not significant, as a significant lack of fit indicates the model does not fit the data within the observed replicate variation and a more complex model has to be considered. R², adjusted R² and predicted R² = 1 indicates perfect adaptation and prediction of the model, therefore the higher these values the better the model. The difference of R²_adj._ and R²_pred._ should be smaller than 0.2, otherwise the model is not correct. The adequate precision, a signal-to-noise ratio, should be greater than 4 to guarantee that the signal is strong enough and can be used for optimization. If curvature appears significant, a quadratic or higher order model is required to model the relationship between the factors and the response with a response surface design (Design Expert, Stat-Ease).

#### 2.6.2. Optimization of screened components

On the basis of the results obtained from the factorial design in the first screening, the media composition was further optimized in a second experiment using a response surface methodology (RSM) (Bezerra et al., 2008). Five factors were detected to enhance the biomass growth during the screening experiments. The three remaining factors were kept at a constant level, 50 % of the concentration of the standard Sphagnum medium. A factorial, central composite design (CCD) for five factors was used with replicates at the center points. Each factor was used at five coded levels: −∞, −1, 0, +1, +∞ (Table 2). Code levels −/+1 represent the factorial points as 50 and 100 % of the concentration of the standard Sphagnum medium, 0 the center points and −/+∞ represents the axial points on the axis of the design space with a defined distance (α) set at 1.49535 coded units from the design center. All linear and interaction terms can be calculated by the facorial points. The axial points can be used for estimation of the quadratic terms.

**Table 2:** The factors and their levels for central composite design used for media optimization towards biomass production of *S. fuscum*, *S. palustre* and *S. squarrosum*. The first column gives the Sphagnum species with their individual medium components without previous statistical significance (50 % of the standard Sphagnum medium). Code levels −/+∞ represents the axial points, −/+ 1 the factorial points as reduced and elevated concentration and 0 the center point of each factor. The chemical formulas of hydrated salts are expressed without water molecules.

| Symbols | Factor | Code Levels |  |  |  |  |
| --- | --- | --- | --- | --- | --- | --- |
| | | $-\infty$ | -1 | 0 | +1 | $+\infty$ |
| *S. fuscum* |  |  |  |  |  |  |
| A | Sucrose (g L<sup>-1</sup>) | 1.28 | 5 | 12.5 | 20 | 23.72 |
| B | NH<sub>4</sub>NO<sub>3</sub> (mM) | 0.47 | 0.63 | 0.94 | 1.25 | 1.40 |
| C | KH<sub>2</sub>PO<sub>4</sub> (mM) | 0.69 | 0.92 | 1.38 | 1.84 | 2.07 |
| D | Ca(NO<sub>3</sub>)<sub>2</sub> (mM) | 1.59 | 2.12 | 3.18 | 4.24 | 4.77 |
| E | ME (%) | 37.6 | 50 | 75 | 100 | 123.8 |
| with 1.68 mM KCl, 0.51 mM MgSO<sub>4</sub>, 22.5 μM FeSO<sub>4</sub> |  |  |  |  |  |  |
| *S. palustre* |  |  |  |  |  |  |
| A | Sucrose (g L<sup>-1</sup>) | 1.28 | 5 | 12.5 | 20 | 23.72 |
| B | NH<sub>4</sub>NO<sub>3</sub> (mM) | 0.47 | 0.63 | 0.94 | 1.25 | 1.40 |
| C | KH<sub>2</sub>PO<sub>4</sub> (mM) | 0.69 | 0.92 | 1.38 | 1.84 | 2.07 |
| D | MgSO<sub>4</sub> (mM) | 0.38 | 0.51 | 0.76 | 1.01 | 1.14 |
| E | Ca(NO<sub>3</sub>)<sub>2</sub> (mM) | 1.59 | 2.12 | 3.18 | 4.24 | 4.77 |
| with 1.68 mM KCl, 50 % ME, 22.5 μM FeSO<sub>4</sub> |  |  |  |  |  |  |
| *S. squarrosum* |  |  |  |  |  |  |
| A | Sucrose (g L<sup>-1</sup>) | 1.28 | 5 | 12.5 | 20 | 23.72 |
| B | NH<sub>4</sub>NO<sub>3</sub> (mM) | 0.47 | 0.63 | 0.94 | 1.25 | 1.40 |
| C | Ca(NO<sub>3</sub>)<sub>2</sub> (mM) | 1.59 | 2.12 | 3.18 | 4.24 | 4.77 |
| D | ME (%) | 37.6 | 50 | 75 | 100 | 112.4 |
| E | FeSO<sub>4</sub> (μM) | 16.9 | 22.5 | 33.8 | 45 | 50.6 |
| with 0.92 mM KH<sub>2</sub>PO<sub>4</sub>, 1.68 mM KCl, 0.51 mM MgSO<sub>4</sub> |  |  |  |  |  |  |

The CCD contained a total of 50 experiments that included 32 trials for factorial design, 10 trials for axial points (two for each variable) and eight trials for replication of the center points. The flasks were filled with 200 mL of the required medium, inoculated with 250 mg of moss material and cultivated for four weeks (*S. fuscum* and *S. palustre*) under standard cultivation conditions.

The cultivation of *S. squarrosum* was prolonged for one week, because of the weak growth performance compared to the other two species. Due to space limitations on the rotary shakers, the flasks were not shaken in week one and three and shaken continuously in week two and four (*S. fuscum* and *S. palustre*), and additionally in week five (*S. squarrosum*). The biomass yield was determined at the end of cultivation by measuring the dry weight, and the model was analyzed.

The highest order polynomial model was selected where the sequential p-value ≤ 0.05 is significant, the lack of fit p-value > 0.05 is not significant, adjusted R² and predicted R² are as high as possible, and the model is not aliased. The significant factors had to be selected by backwards elimination of non-significant factors with a p-value > 0.05 or a linear effect has to be selected to support hierarchy. The selected model including the selected factors was checked using ANOVA. Transformation was applied to meet statistical assumptions like normal distribution and a new model has to be selected and checked again as described before (Design Expert, Stat-Ease).

#### 2.6.3. Validation of the optimized media composition determined by the model

The model analyzes the optimized media composition towards biomass production and these predictions need validation. The validation experiment was carried out in the same way as the CCD experiment. Shake flasks experiments (n=3, with exception of the predicted optimized media of *S. squarrosum* n=2) were carried out and the biomass yield between the medium composition of the center points of the CCD and the predicted optimized media concentration were compared.

### 2.7. Statistical analysis

The statistical software package Design-Expert® (Stat-Ease, 11.1.2.0, Minneapolis, USA) was used for regression analysis of experimental data and to plot response surface. ANOVA was used to estimate the statistical parameters.

## 3. Results and Discussion

### 3.1. Screening of important medium components for biomass production

The standard Sphagnum medium was established by Beike *et al*. (2015) with regard to *S. palustre* productivity, and tested for four other *Sphagnum* species. However, this medium was not optimized for biomass production for the 19 *Sphagnum* species tested by Heck *et al*. (2021). Nevertheless, it is a good basis for peat moss cultivation because it comprises all components needed for moss growth: sucrose, NH_4_NO_3_, KH_2_PO_4_, KCl, MgSO_4_, Ca(NO_3_)_2_, micro elements (ME) and FeSO_4_.

A first screening of the influence of these eight components was important prior to optimization. The treatment of the components as a salt and not as ions as one factor was neccesary to keep the number of trials small. This reduced the number of factors to eight and with it the number of experiments. With fractional factorial designs some limitations can occur for the estimation of the main and interaction effects, because some are estimated together (Candioti et al., 2014). Another limitation relies on the fact that they have only two levels for each input factor, resulting in a linear model (Fukuda et al., 2018). To prove linearity of the model, center points have to be added at the exact mid-point of all factor settings to evaluate curvature and identify significant second-order effects (Bezerra et al., 2008).

The experimental matrix including 16 trials for the 8 different variables and the dry weight as response for each setup is shown in Table S1 for *S. fuscum*, Table S2 for *S. palustre* and Table S3 for *S. squarrosum*, sorted by the standard order (Std), which sorts the factor settings in low to high concentration pattern. The highest tested concentration corresponds to the standard Sphagnum medium and 10 % of this medium to the lowest concentration adjusted for *S. palustre*. This yielded in a biomass amount of 28.55 to 100.64 mg DW with a maximum yield of 95.59 ±5.05 mg DW l^−1^ for Std 31 and 32, representing the standard Sphagnum medium (Table S2). To reduce the wide range of biomass yield, the range between the factors could be narrowed.

As the highest *S. palustre* biomass was gained with the highest tested concentration of all factors, the lower value of all factors was set to 50 % for the screening experiments of *S. fuscum* and *S. squarrosum*. *S. fuscum* grew between 33.35 to 65.55 mg DW L^−1^, where the standard Sphagnum medium yielded the highest biomass (Table S1, Std 31, 32). The biomass of *S. squarrosum* yielded between 10.11 and 45.55 mg DW L^−1^. Compared to the other two species, the standard Sphagnum medium resulted in an only moderate biomass increase for this species, whereas the reduction of NH_4_NO_3_, KCl, MgSO_4_ and ME yielded the highest biomass (Table S3). The ANOVA results are presented in Table 3, where all three models are significant with a non-significant lack of fit, indicating no reason to doubt the fitness of the model. Out of the eight variables studied, the biomass of *S. fuscum* was positively influenced by an increase in the concentration of sucrose, NH_4_NO_3_, KH_2_PO_4_, Ca(NO_3_)_2_ and ME (Table 3). The biomass of *S. palustre* was positively influenced by an increase in the concentration of sucrose, NH_4_NO_3_, KH_2_PO_4_, MgSO_4_ and Ca(NO_3_)_2_ (Table 3). The biomass of *S. squarrosum* was positively influenced by an increase in the concentration of sucrose and Ca(NO_3_)_2_, and by a decrease in the concentration of NH_4_NO_3_, ME and FeSO_4_ (Table 3). Especially as the significant curvature test of *S. palustre* and *S. squarrosum* indicates non-linearity of the model (Table 3), a more complex model has to be used to identify the optimized media concentration.

**Table 3:**
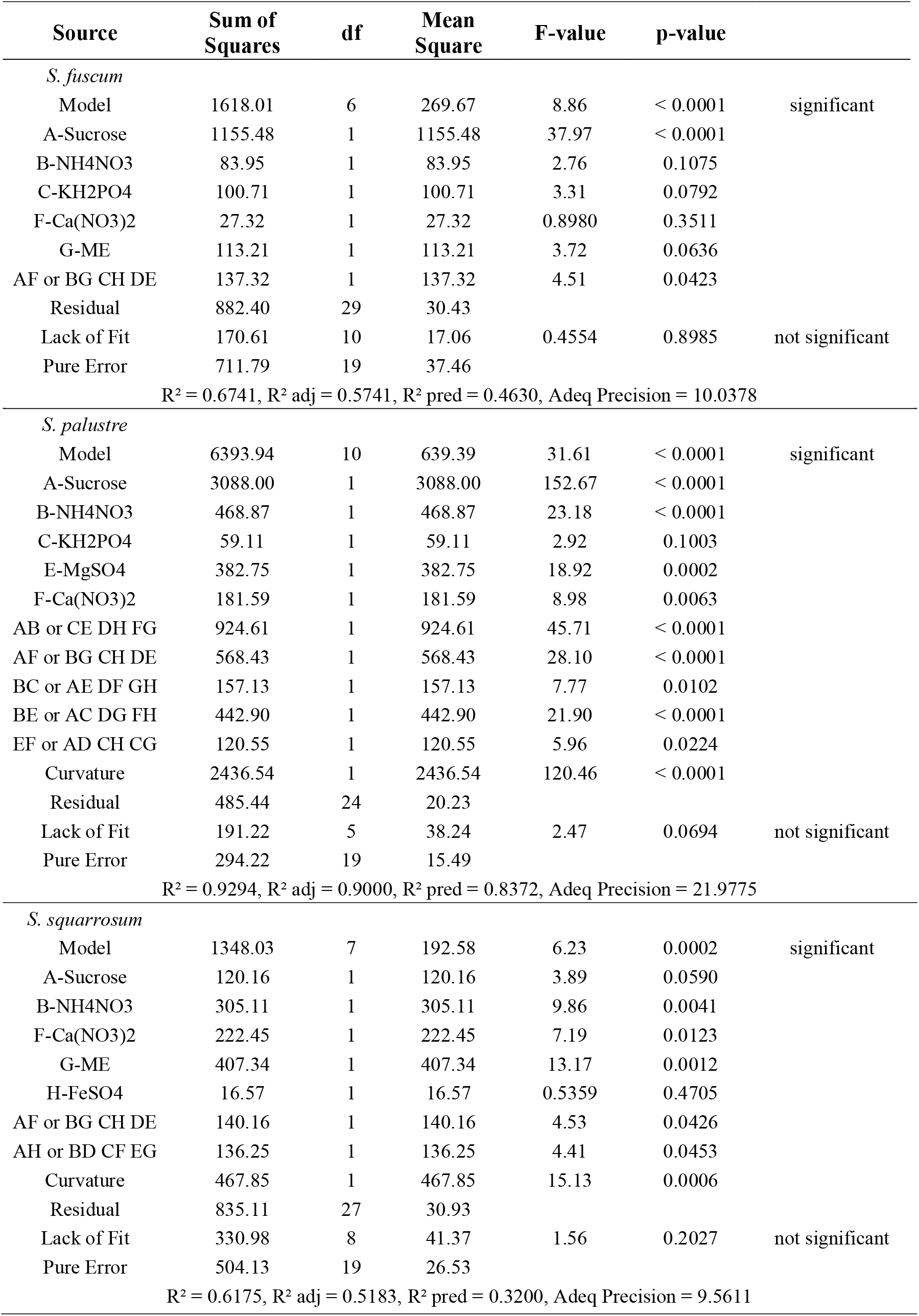
Analysis of variance (ANOVA) generated by Design Expert software (Stat-Ease, 11.1.2.0) for the two-level factorial design used as first screening for the optimized media composition of S. fuscum, S. palustre and S. squarrosum in relation to biomass production. Sum of squares, degree of freedom (df), mean square, F-value and P-value. R-squared (R^2^), adjusted R-squared (R^2^ adj), predicted R-squared (R^2^ pred) and Adequate Precision (Adeq Precision).

In this two-level factorial design, the highest *S. palustre* biomass yield was achieved with the standard Sphagnum medium with 2 % sucrose. The second highest *S. fuscum* biomass yield was also achieved with that medium. In contrast, the productivity of *S. squarrosum* was on an average with that medium. This correlates with the findings of Heck et al. (2021), who reported that the nutrient composition of the standard Sphagnum medium is suboptimal for some *Sphagnum* species and has to be improved further.

The productivity of all three *Sphagnum* species was positively influenced by sucrose. These growth-promoting effects of sugar have also been found in previous *in-vitro* cultivation studies (Beike et al., 2015; Rudolph et al., 1988; Simola, 1969). *Sphagnum* is able to take up and use exogenous organic compounds, exhibiting a mixotrophic lifestyle (Graham et al., 2010) with sucrose as one of the best carbon sources (Rudolph et al., 1988; Simola, 1969).

The nitrogen sources NH_4_NO_3_ and Ca(NO_3_)_2_ also positively influence the productivity of all three *Sphagnum* species. This is in accordance with the positive effect of 1.25 mM NH_4_NO_3_ on the growth of *S. nemoreum* (Simola, 1975), and combined with sucrose on the growth of *S. palustre* (Beike et al., 2015). The effect of Ca(NO_3_)_2_ cannot be attributed to one of the ions due to ion confounding. Ion confounding occurs through the use of salts, because changing the concentration of a single cation or anion results in a simultaneous change in the associated co-ion (Niedz and Evens, 2006). Varying Ca(NO_3_)_2_ varies both the Ca^2+^ and NO_3_^−^ ions simultaneously. Any change in the output may be due to the varied ion concentration of Ca^2+^ or NO_3_^−^ or the interaction between Ca^2+^ and NO_3_^−^. The use of salts instead of ions as a factor impaired the detailed understanding of the metabolism as the ion-specific effects are not obvious. Nevertheless, the optimization of the media components was the main focus of this study and the growth-influencing factors were identified.

### 3.2. Optimization of the screened medium

Once the screening process identified the relevant factors and discarded the insignificant ones, a more complex response surface is required (Fukuda et al., 2018). To model a second order response surface, optimization designs use three to five levels of each input factors, which increases the number of required experiments. One of the most often used optimization designs is the central composite design (CCD), where all factors are studied in five levels (Bezerra et al., 2008).

The variables in the two-level factorial design showing a significant effect on the biomass production were selected, and the remaining factors were set at 50 % of the standard Sphagnum medium: 0.92 mM KH_2_PO_4_, 1.68 mM KCl, 0.51 mM MgSO_4_, 50 % ME and 22.5 µM FeSO_4_. The experimental design and results obtained for biomass production of the CCD are shown in Table S4 for *S. fuscum*, Table S5 for *S. palustre* and Table S6 for *S. squarrosum*.

The best fitting model was a reduced quadratic response surface including inverse square root transformation for *S. fuscum*, whilst the data for *S. palustre* and *S. squarrosum* did not require transformations to fit statistical assumptions. In the case of *S. fuscum*, sucrose and Ca(NO_3_)_2_ as well as the quadratic effects of sucrose, NH_4_NO_3_ and KH_2_PO_4_ had a positive effect on biomass production. The productivity of *S. palustre* was positively influenced by linear effects of sucrose and NH_4_NO_3_ as well as by the quadratic effects of NH_4_NO_3_, MgSO_4_ and Ca(NO_3_)_2_. Sucrose and NH_4_NO_3_ had the same positive effect on the biomass production of *S. squarrosum* as well as interaction effects of sucrose with NH_4_NO_3_ and ME with FeSO_4_, and quadratic effects of sucrose (Table 4).

**Table 4:** Analysis of variance (ANOVA) generated by Design Expert software (Stat-Ease, 11.1.2.0) of the quadratic models for the central composite design used as opimization of the media composition of *S. fuscum* including inverse square root transformation, *S. palustre* and *S. squarrosum* in relation to biomass production. Sum of squares, degree of freedom (df), mean square, F-value and P-value. R-squared (R^2^), adjusted R-squared (R^2^ adj), predicted R-squared (R^2^ pred) and Adequate Precision (Adeq Precision).

| Source | Sum of Squares | df | Mean Square | F-value | p-value |  |
| --- | --- | --- | --- | --- | --- | --- |
| *S. fuscum* |  |  |  |  |  |  |
| Model | 0.0011 | 7 | 0.0002 | 9.28 | < 0.0001 | significant |
| A-Sucrose | 0.0004 | 1 | 0.0004 | 22.45 | < 0.0001 |  |
| B-NH4NO3 | 0.0000 | 1 | 0.0000 | 0.8903 | 0.3508 |  |
| C-KH2PO4 | 0.0000 | 1 | 0.0000 | 1.48 | 0.2303 |  |
| D-Ca(NO3)2 | 0.0001 | 1 | 0.0001 | 3.27 | 0.0776 |  |
| A<sup>2</sup> | 0.0002 | 1 | 0.0002 | 11.84 | 0.0013 |  |
| B<sup>2</sup> | 0.0001 | 1 | 0.0001 | 3.64 | 0.0631 |  |
| C<sup>2</sup> | 0.0001 | 1 | 0.0001 | 3.27 | 0.0776 |  |
| Residual | 0.0007 | 42 | 0.0000 |  |  |  |
| Lack of Fit | 0.0007 | 35 | 0.0000 | 6.41 | 0.0080 | not significant |
| Pure Error | 0.0000 | 7 | 3.018E-06 |  |  |  |
| $R^2 = 0.6072$, $R^2$ adj = 0.5418, $R^2$ pred = 0.3505, Adeq Precision = 9.7669 | | | | | | |
| *S. palustre* |  |  |  |  |  |  |
| Model | 1.046E+05 | 7 | 14936.59 | 7.95 | < 0.0001 | significant |
| A-Sucrose | 33776.79 | 1 | 33776.79 | 17.97 | 0.0001 |  |
| B-NH4NO3 | 6072.73 | 1 | 6072.73 | 3.23 | 0.0794 |  |
| D-MgSO4 | 579.41 | 1 | 579.41 | 0.3083 | 0.5817 |  |
| E-Ca(NO3)2 | 216.48 | 1 | 216.48 | 0.1152 | 0.7360 |  |
| B<sup>2</sup> | 6204.97 | 1 | 6204.97 | 3.30 | 0.0764 |  |
| D<sup>2</sup> | 6419.33 | 1 | 6419.33 | 3.42 | 0.0716 |  |
| E<sup>2</sup> | 19632.16 | 1 | 19632.16 | 10.45 | 0.0024 |  |
| Residual | 78934.78 | 42 | 1879.40 |  |  |  |
| Lack of Fit | 73014.84 | 35 | 2086.14 | 2.47 | 0.1075 | not significant |
| Pure Error | 5919.94 | 7 | 845.71 |  |  |  |
| $R^2 = 0.5698$, $R^2$ adj = 0.4981, $R^2$ pred = 0.3063, Adeq Precision = 10.3522 | | | | | | |
| *S. squarrosom* |  |  |  |  |  |  |
| Model | 81613.04 | 7 | 11659.01 | 8.69 | < 0.0001 | significant |
| A-Sucrose | 42609.81 | 1 | 42609.81 | 31.74 | < 0.0001 |  |
| B-NH4NO3 | 11690.92 | 1 | 11690.92 | 8.71 | 0.0052 |  |
| D-ME | 5871.59 | 1 | 5871.59 | 4.37 | 0.0426 |  |
| E-FeSO4 | 5339.73 | 1 | 5339.73 | 3.98 | 0.0526 |  |
| AB | 7919.11 | 1 | 7919.11 | 5.90 | 0.0195 |  |
| DE | 4213.62 | 1 | 4213.62 | 3.14 | 0.0837 |  |
| A<sup>2</sup> | 3968.26 | 1 | 3968.26 | 2.96 | 0.0929 |  |
| Residual | 56377.14 | 42 | 1342.31 |  |  |  |
| Lack of Fit | 48734.03 | 35 | 1392.40 | 1.28 | 0.3950 | not significant |
| Pure Error | 7643.11 | 7 | 1091.87 |  |  |  |
| $R^2 = 0.5914$, $R^2$ adj = 0.5233, $R^2$ pred = 0.4260, Adeq Precision = 11.9436 | | | | | | |

Three-dimensional response surfaces were plotted on the basis of the final model, in order to investigate the interaction between two variables on the biomass production, whereby the remaining variables were kept constant at their optimum level. The interaction between NH_4_NO_3_ and sucrose shows the nonlinear effect of these factors on the biomass production of *S. fuscum*, *S. palustre* and *S. squarrosum* (Figure 1). The optimal concentration may lie outside the ranges initially chosen for *S. palustre* and *S. squarrosum*, as the highest predicted biomass is at the border. However, increased sucrose concentrations of 5% or higher negatively affected peat moss productivity (Beike et al., 2015; Simola, 1969).

**Figure 1:**
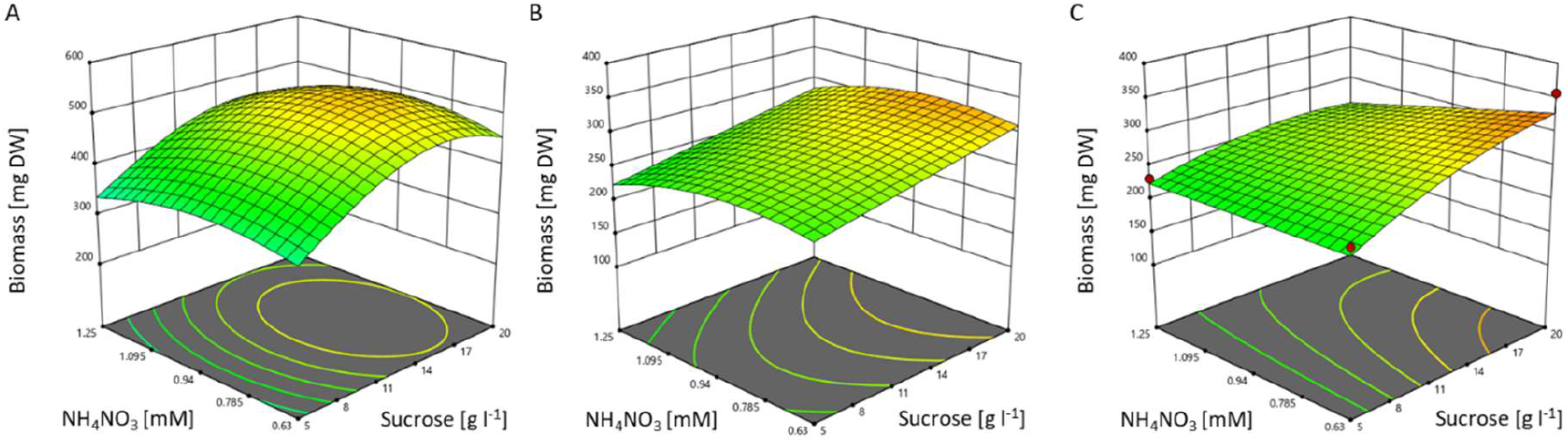
3D response surface for biomass production of *S. fuscum*, *S. palustre* and *S. squarrosum*. The plot shows the effect of interaction between NH4NO3 and sucrose of A) *S. fuscum* (KH2PO4, Ca(NO3)2 and ME were kept konstant at 1.29 mM, 2.12. mM and 50%, respectively), B) *S. palustre* (KH2PO4, MgSO4 and Ca(NO3)2 were kept konstant at 0.92 mM, 0.78. mM and 3.14 mM, respectively) and C) *S. squarrosum* (Ca(NO3)2, ME and FeSO4 were kept konstant at 2.12 mM, 50% and 22.5 µM, respectively).

In the CCD experiments, *S. squarrosum* yielded the highest biomass with Std 2 (Table S6), which is in agreement with the model prediction that the optimized medium composition has a high concentration of sucrose and low concentrations of the remaining nutrients (Table 5). *S. fuscum* and *S. palustre* yielded the highest biomass in Std 47 (Table S4) and Std 46 (Table S5), both representing one center point. This corresponds to the model prediction using the Design Expert Software, where the optimal nutrient concentration is similar to the medium composition at the center points of *S. fuscum* and *S. palustre* (Table 5).

**Table 5:** Optimized media composition of *S. fuscum*, *S. palustre* and *S. squarrosum*. The amount of nutrients in 100 % of the standard Sphagnum medium is compared with the optimized fuscum, palustre and squarrosum media. The chemical formulas of hydrated salts are expressed without water molecules.

|  | **Standard**<br>**Sphagnum**<br>**Medium (100 %)** | *S. fuscum* | *S. palustre* | *S. squarrosus* |
| --- | --- | --- | --- | --- |
| Sucrose | 20 (g L<sup>-1</sup>) | 80 % | 100 % | 100 % |
| NH<sub>4</sub>NO<sub>3</sub> | 1.25 (mM) | 71 % | 68 % | 50 % |
| KH<sub>2</sub>PO<sub>4</sub> | 1.84 (mM) | 70 % | 50 % | 50 % |
| KCl | 3.35 (mM) | 50 % | 50 % | 50 % |
| MgSO<sub>4</sub> | 1.01 (mM) | 50 % | 77 % | 50 % |
| Ca(NO<sub>3</sub>)<sub>2</sub> | 4.24 (mM) | 50 % | 74 % | 50 % |
| ME | 100 % | 50 % | 50 % | 50 % |
| FeSO<sub>4</sub> | 45 (μM) | 50 % | 50 % | 50 % |

### 3.3. Validation of the optimized medium

To verify the obtained optimized media concentrations with regard to improved biomass yields, validation experiments were conducted. All three validation experiments yielded higher biomasses by using the optimized media concentration as specified by the Design Expert Software. For *S. fuscum*, a biomass of 491.6 ± 24.7 mg DW was obtained by using the optimized concentrations, compared to 403.4 ± 5.8 mg DW by using the concentrations of the center point, which yielded the highest biomass during the optimization experiment. The optimized media of *S. palustre* yielded a maximum biomass of 473.6 ± 10.9 mg DW compared to 402.0 ± 24.8 mg DW by using the concentrations of the center point representing the best productivity during the optimization experiment. *S. squarrosum* yielded 707.6 ± 5.3 mg DW in the validation experiment with the predicted optimized media, whereas the same media composition yielded only 356.8 mg DW during the CCD (Table S6, Std 2).

This variance in biomass production is most likely a consequence of heterogenous starting material. The inocula of all three species were treated in the same way, but the preculture could differ concerning length of the gametophores and number of capitula, because the moss material was disrupted manually with forceps. Vegetative growth of peat mosses is possible from several parts of the shoot, like capitula, fascicles, branches and stems (Poschlod & Pfadenhauer, 1989). Green stems and apical branches showed the highest regeneration potential for *S. palustre*, while brown parts and leaves did not regenerate (Sobotka, 2015), and the regeneration potential of *S. angustifolium* capitula was up to ten times higher than out of stems (Tuittila et al., 2003), which is in line with our own observations. The inocula may vary in the composition of parts of the gametophores and therefore may have a varying regeneration potential. The use of a defined number of capitula as inoculum may thus result in a more homogenous batch-to-batch productivity. However, such a labour-intensive procedure is not appropriate for the production of founder material for Sphagnum farming.

The concentrations of KCl, ME and FeSO_4_ are set at 50 % of the standard Sphagnum medium in the optimized media composition of *S. fuscum*, *S. palustre* and *S. squarrosum*, whereas the concentration of sucrose, NH_4_NO_3_, KH_2_PO_4_, MgSO_4_ and Ca(NO_3_)_2_ varies between all three species, which may reflect the nutrition status of their respective habitats: *S. fuscum* is an ombrotrophic (rain-fed) species and grows predominantly in nutrient-poor oligotrophic and mesotrophic mires; *S. palustre* grows in a wide range of mesotrophic peatlands containing intermediate levels of nutrients and is absent only from strongly acidic locations; *S. squarrosum* grows predominantly in mesotrophic to slightly eutrophic areas (Daniels and Eddy, 1990). Although *S. squarrsoum* can be found in the most nutrient-rich locations, the optimized medium has the lowest nutrient concentrations with the exception of sucrose. *S. palustre* needed the same concentration of sucrose and the highest concentrations of MgSO_4_ and Ca(NO_3_)_2_ compared to the other two species. In contrast, the nutrient-poor adapted *S. fuscum* needed the lowest concentration of sucrose as well as the highest concentrations of NH_4_NO_3_ and KH_2_PO_4_. The phosphate mobility of oligotrophic raised-bog soils is higher than that of mineral soils (Kuntze and Scheffer, 1979), which could explain the higher consumption of PO_4_^3-^.

Sucrose concentration played a significant role for peat moss productivity in all three optimized media. This is in agreement with the literature: The growth of *S. nemoreum* could be increased by addition of sucrose, glucose, fructose and mannose with 1 % sucrose as the best carbon source (Simola, 1969). Also in Beike et al. (2015), 2 % sucrose significantly increased the productivity of *S. palustre* compared to 0.3 % sucrose. In contrast, *S. imbricatum* utilized glucose as the main carbon and energy source for their growth (Kajita et al., 1987).

In nature, submerged tissues may be limited in photosynthesis by the availability of dissolved organic carbon and/or light (Graham et al., 2010), wheras emergent tissues are less carbon-limited, as CO_2_ diffuses around 1000 times faster in air than in water (Proctor et al., 1992). Under wet conditions hummock-forming species can also be covered with a water layer and may be carbon-limited (Smolders et al., 2001). Glucose and other sugars occur in the peat and peat water from decomposing organic matter, or are exuded from the roots of nearby vascular plants (Graham et al., 2010). The uptake of sugars (mixotrophy) helps peat mosses to deal with carbon limitations (Graham et al., 2010).

*S. fuscum* forms compact hummocks on raised or blanket mires, while *S. palustre* and *S. squarrosum* can be found in wet woodlands, ditches and flushes (Atherton et al., 2010). This correlates with the higher required amount of sucrose for the submerged species *S. palustre* and *S. squarrosum* as compared to the emergent species *S. fuscum*. However, at current knowledge an optimal medium composition in the laboratory can not be predicted from the knowledge of the natural habitat. Besides the multitude of inorganic salts and sugars described here, specific microbiome compositions in the natural habitat (e.g., Holland-Moritz et al., 2021) may explain different nutrient requirements between the field and the axenic laboratory culture.

### 3.4. Cultivation in the photobioreactor

The optimization of media composition was validated on a larger scale in 5 L photobioreactors by comparing productivities in the standard and the optimized medium. A direct comparison of the biomass yield between standard and optimized medium is only possible when the cultivation periods are identical. Due to the different duration of the cultivation in both media, the average growth rate in g·d^−1^ was used (Figure 2).

**Figure 2:**
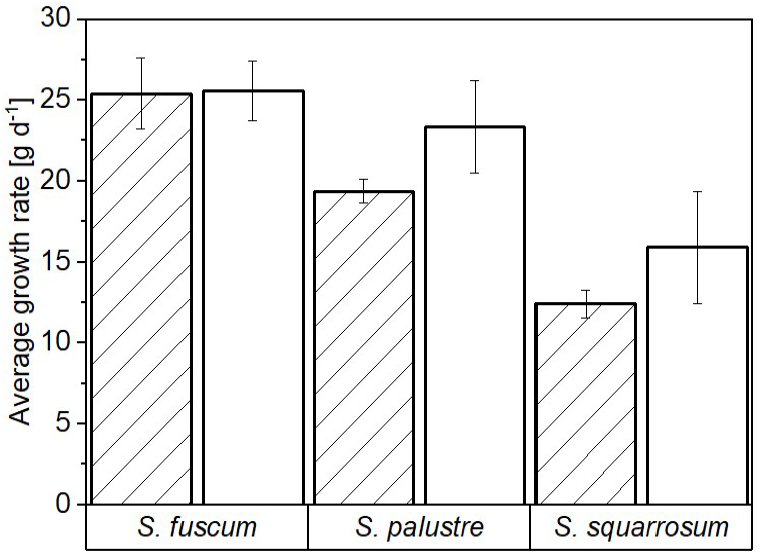
Average growth rates of *S. fuscum*, *S. palustre* and *S. squarrosum* in 5 L photobioreactors. The *Sphagnum* species were cultivated in (hatched bar) standard Sphagnum medium (n=2), where the cultivation period varies between 23 and 28 days and in (empty bar) optimized medium (n=3), where the bioreactor run was set to 24 days.

*S. fuscum* had similar growth rates between the two tested media, but in the optimized medium needed less nutrients, which is economically sensible. The optimized medium of *S. palustre* and *S. squarrosum* not only needed less nutrients, it also increased productivity. Nevertheless, *S. squarrosum* showed the lowest average growth rate in the bioreactor among the three species. This is in contrast to the axenic *in-vitro* cultivation in flasks, where *S. squarrosum* yielded the highest biomass compared to *S. fuscum* and *S. palustre* in the standard Sphagnum medium (Heck et al., 2021). One possible explanation for this discrepancy is the difference in the cultivation technology; rotating flasks versus stirred bioreactors. Consequently, experiments on shear-stress sensitivity of this peat moss species may show in future whether different hydrodynamic and mixing conditions in the bioreactor can impede the productivity of *S. squarrosum*.

Cultivation of about 15 g start FW in the photobioreactor containing 5 L of the respective optimized medium for 24 days resulted in 628.67 ± 36.67 g FW of *S. fuscum*, 576.33 ± 6.24 g FW of *S. palustre* and 398.50 ± 31.83 g FW of *S. squarrosum* (Table 6) with a fresh to dry weight ratio of 9.2 ± 0.3 (n=9). This is in contrast to Beike et al. (2015), where the ratio of fresh to dry weight is approximately 14 ± 2.7 (n=12) for *S. palustre*. This variance is a consequence of the different methods of fresh weight determination. In both cases the moss material was filtered for 1 minute, but in this study, closing the funnel and generating a vacuum removes more water, which leads to a lower and less varying fresh to dry weight ratio.

**Table 6:** Average growth rates of *S. fuscum*, *S. palustre* and *S. squarrosum* in 5 L photobioreactors. The *Sphagnum* species were cultivated in standard Sphagnum medium (n=2), where the cultivation period varies between 24 and 28 days and in optimized medium (n=3), where the bioreactor run was set to 24 days.

| Media | Cultivation Time<br>d | Inocula |  | Harvest |  | Average Growth Rate |  |
| --- | --- | --- | --- | --- | --- | --- | --- |
|  |  | g FW | g DW | g FW | g DW | g FW/d | g DW/d |
| *S. fuscum* |  |  |  |  |  |  |  |
| Standard | 28 |  |  | 726.05 ± 61.45 |  | 25.41 ± 2.19 |  |
| Optimized | 24 | 14.53 ± 0.19 | 1.52 ± 0.03 | 628.67 ± 36.67 | 68.63 ± 1.85 | 25.59 ± 1.85 | 2.79 ± 0.08 |
| *S. palustre* |  |  |  |  |  |  |  |
| Standard | 24 |  |  | 480.85 ± 17.15 |  | 19.38 ± 0.71 |  |
| Optimized | 24 | 15.63 ± 0.19 | 1.75 ± 0.02 | 576.33 ± 6.24 | 64.63 ± 2.84 | 23.36 ± 2.84 | 2.62 ± 0.12 |
| *S. squarrosum* |  |  |  |  |  |  |  |
| Standard | 28 |  |  | 363.85 ± 23.85 |  | 12.40 ± 0.85 |  |
| Optimized | 24 | 16.63 ± 0.50 | 2.32 ± 0.07 | 398.50 ± 31.83 | 42.37 ± 3.46 | 15.91 ± 3.46 | 1.67 ± 0.14 |

Compared to the cultivation of *S. palustre* in the photobioreactor of Beike et al. (2015), precultivation of the inocula, aeration of the bioreactor with 2 % CO_2_ and optimization of the standard Sphagnum medium shortened the cultivation time from about 30 days to 24 days and furthermore increased the biomass production from 30-fold to nearly 40-fold in our study.

The optical assessment of biomass increase is depicted in Figure 3 for *S. fuscum*, Figure 4 for *S. palustre* and Figure 5 for *S. squarrosum*. In the first week of cultivation the biomass amount in the bioreactor remained constant. This might be connected to the lag phase that Beike et al. (2015) reported. Visually, an increase in biomass was evident from the images taken on day 7. From day 13 to 17, depending on the species, the bioreactor was filled with the produced biomass. There were still some free spaces inbetween the gametophores for growth, which were filled at the end of the cultivation. At first the moss was bright green and became darker and partly brownish towards the end (Figure 2-4, A), which seems to have no effect on the vitality of the moss. This color change may be due to high light availability. In nature, the majority of the species are green when shaded and develop secondary pigments when well-illuminated (Atherton et al., 2010). *S. fuscum* is found to be mid to deep brown and rarely all green. *S. palustre* is pale green or yellow-brown and occasionally the whole plant is green, whereas *S. squarrosum* varies from pale green to yellow-green and rarely pale brown (Daniels and Eddy, 1990).

**Figure 3:**
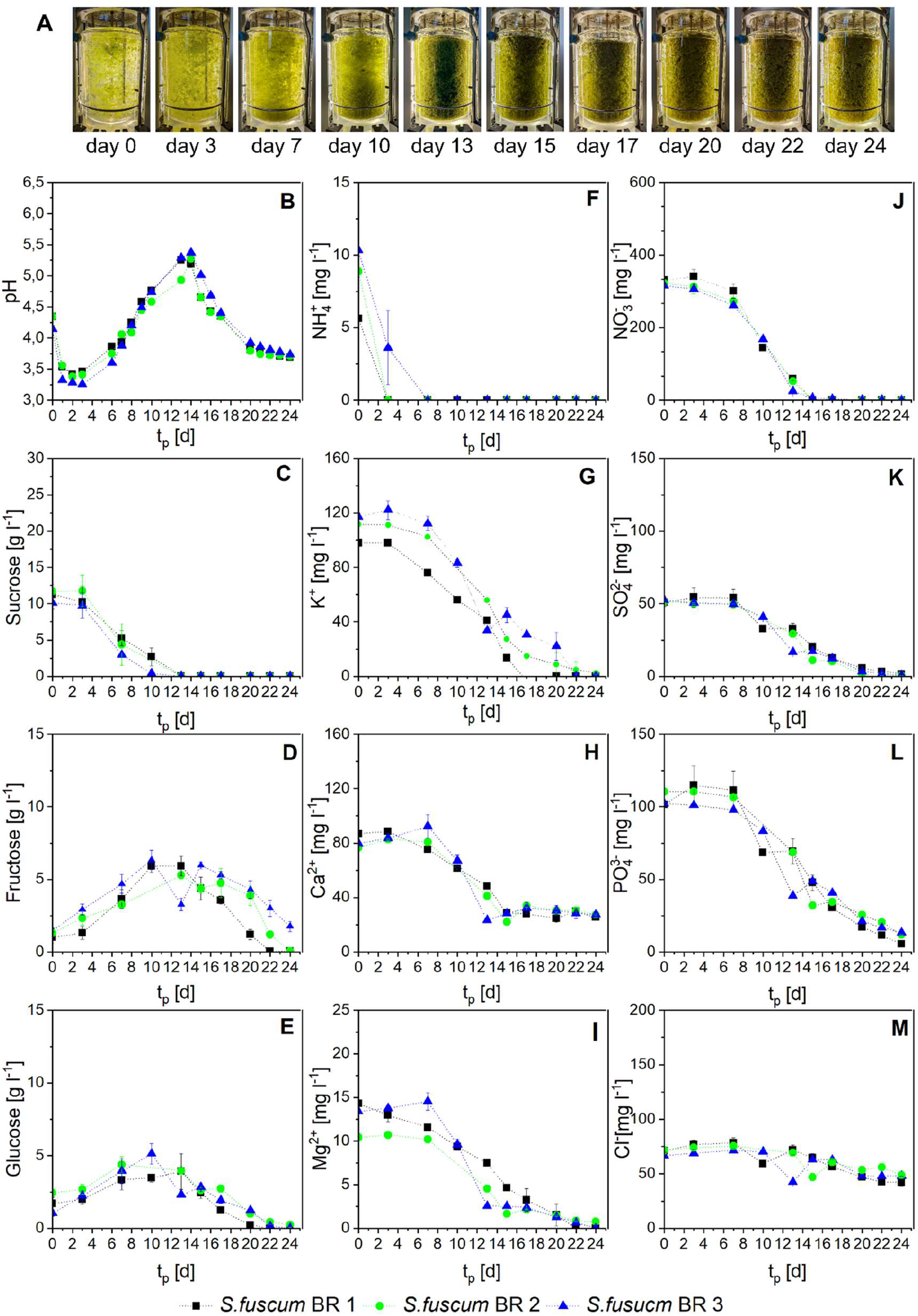
Cultivation of *S. fuscum* in the bioreactor. Biomass increase of *S. fuscum* documented photographically, pH changes and nutrient concentrations during bioreactor cultivation in optimized Fuscum medium. A) Pictures were taken directly after inoculation and 3, 7, 10, 13, 15, 17, 20, 22 and 24 days thereafter. The y-axis shows: B) pH changes during the bioreactor cultivation, the concentration of C) sucrose, D) fructose and E) glucose in mg per liter, the cation concentrations F) NH4^+^, G) K^+^, H) Ca^2+^, I) Mg^2+^ in mg per liter and the anion concentrations of J) NO3^−^, K) SO4^2-^, L) PO4^3-^, M) Cl^−^ in mg per liter, while the x-axis shows the day of cultivation.

**Figure 4:**
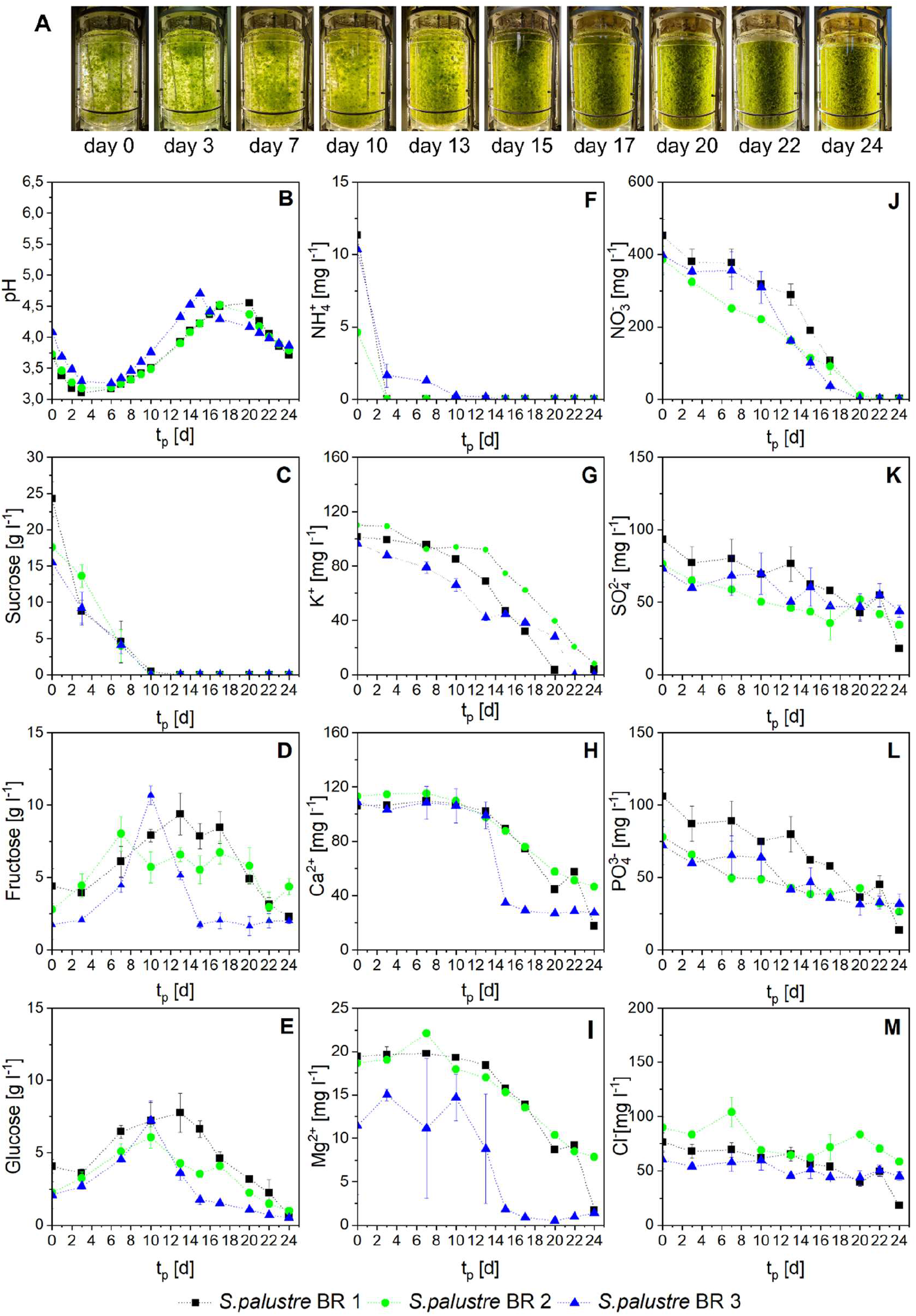
Cultivation of *S. palustre* in the bioreactor. Biomass increase of *S. palustre* documented photographically, pH changes and nutrient concentrations during bioreactor cultivation in optimized Palustre medium. A) Pictures were taken directly after inoculation and 3, 7, 10, 13, 15, 17, 20, 22 and 24 days thereafter. The y-axis shows: B) pH changes during the bioreactor cultivation, the concentration of C) sucrose, D) fructose and E) glucose in mg per liter, the cation concentrations concentrations F) NH4^+^, G) K^+^, H) Ca^2+^, I) Mg^2+^ in mg per liter and the anion concentrations of J) NO3^−^, K) SO4^2-^, L) PO4^3-^, M) Cl^−^ in mg per liter, while the x-axis shows the day of cultivation.

**Figure 5:**
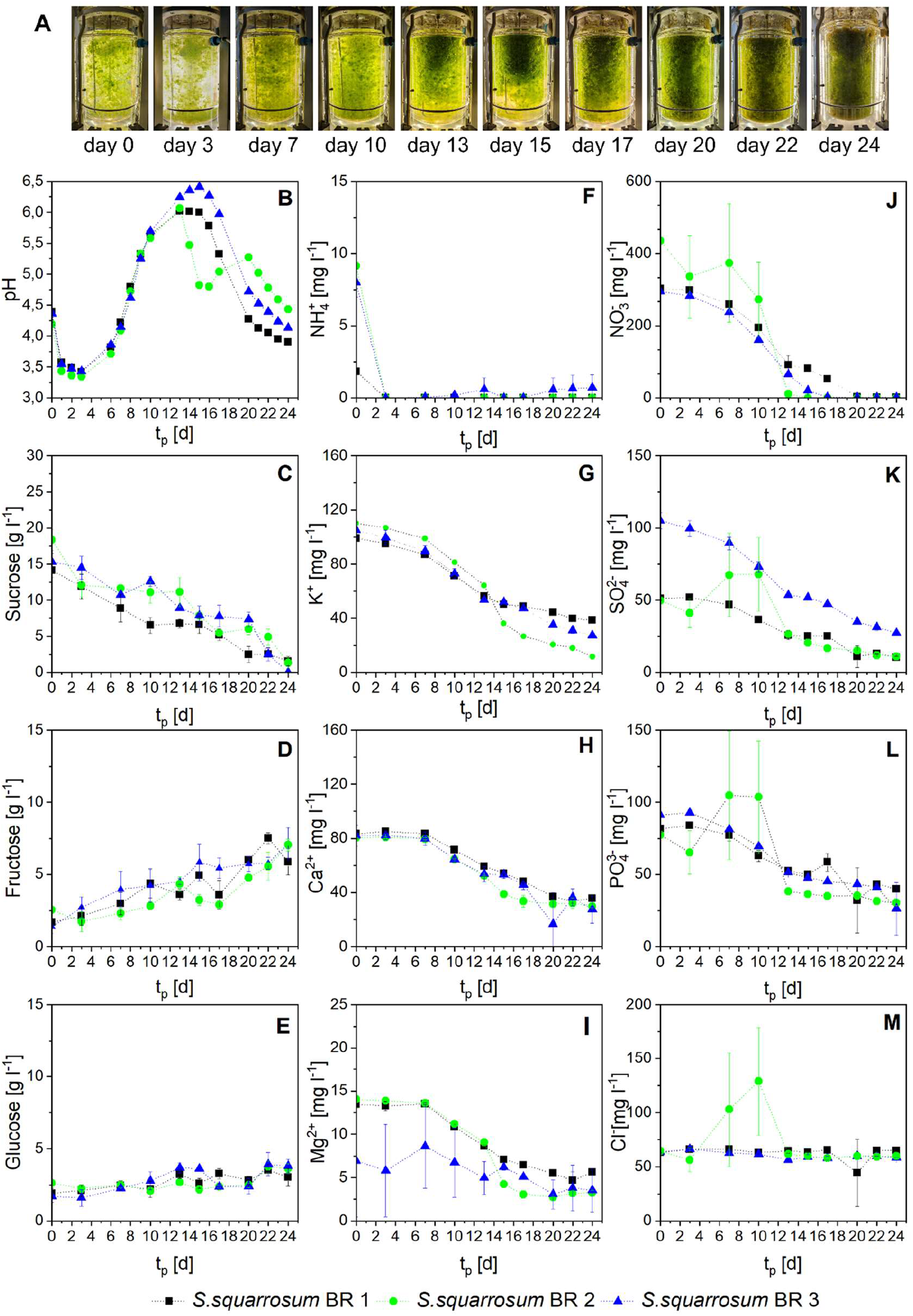
Cultivation of *S. squarrosum* in the bioreactor. Biomass increase of *S. squarrosum* documented photographically, pH changes and nutrient concentrations during bioreactor cultivation in optimized Squarrosum medium. A) Pictures were taken directly after inoculation and 3, 7, 10, 13, 15, 17, 20, 22 and 24 days thereafter. The y-axis shows: B) pH changes during the bioreactor cultivation, the concentration of C) sucrose, D) fructose and E) glucose in mg per liter, the cation concentrations concentrations F) NH ^+^, G) K^+^, H) Ca^2+^, I) Mg^2+^ in mg per liter and the anion concentrations of J) NO3^−^, K) SO4^2-^, L) PO4^2-^, M) Cl^−^ in mg per liter, while the x-axis shows the day of cultivation.

The pH was not adjusted during cultivation according to the findings of Beike et al. (2015), a fixed pH was not suitable for the cultivation of *S. palustre* in the photobioreactor. We also observed the pH drop after autoclaving caused by precipitation of some nutrients (Beike et al., 2015). In addition, the inoculation with a two-week-old precultre decreased the starting pH further. The changes in pH during the bioreactor cultivation of all *Sphagnum* species was similar (Figure 2-5, B). After starting the cultivation, the pH decreased in the first three days from almost pH 4.4 for *S. fuscum* and *S. squarrosum* and an initial pH of 3.83 ± 0.17 for S*. palustre* to nearly pH 3.1. This acidification is related to ion exchange, as cations were taken up by the surface of the plants, a phenomenon observed in nature (Clymo, 1963, 1964) as well as in *in-vitro* cultures (Beike et al., 2015; Rudolph et al., 1988). Especially ammonia uptake correlated with the pH decrease (Figure 3-5, F). After reaching the pH minimum, the whole amount of ammonia is taken up. This result is explained by the release of H^+^ ions through the assmiliation of ammonium ions in the cytoplasm (Kirkby, 1968; Raven and Smith, 1976). During further cultivation, the pH increased again up to pH 5.27 ± 0.08 at day 14 for *S. fuscum*, up to pH 4.7 between day 15 to 20 for *S. palustre* and up to pH 6.41 between day 13 to 15 for *S. squarrosum*, accompanied by the uptake of nitrate ions. The assimilation of nitrate releases OH^−^ ions. To keep the pH value constant in the cytoplasm excess OH^−^ is excreted from the cell (Raven and Smith, 1976). The complete uptake of nitrate is in correlation with the pH maxima (Figure 3-5 B,J). After the nitrate is taken up, the pH started to decrease again to nearly 3.7 for all tested species. It seems that nitrogen deprivation affects the pH value of the medium. Rasmussen et al. (1995) reported about excretion of Sphagnum acid into the culture media of *S. fallax* and *S. cuspidatum*. However, we could not find any reported correlation between nitrogen starvation and Sphagnum acid production.

The analysis of the nutrient concentrations in the medium (Figure 3-5, F,J) revealed a rapid decrease of ammonia during the first three cultivation days, while nitrate was taken up more slowly. It is obvious that ammonia is the preferred nitrogen source in the three species tested here, which is in agreement with data about *S. nemoreum* (Simola, 1975). During all bioreactor runs we observed that nitrate was completely depleted from the medium before the end of the experiment. Nevertheless, the results of DOE showed that higher concentration of nitrate did not lead to higher biomass yield. Higher light availabilty and additional CO_2_ supplementation in bioreactor cultures increase the photosynthetic acivity of *Sphagnum* (Haraguchi and Yamada, 2011; Jauhiainen and Silvola, 1999). This could result in better growth responses with higher nutrient requirements compared to flask cultivation, which were conducted without supplementation of CO_2_.

The optimized media contained 1.6 % sucrose for *S. fuscum* and 2 % sucrose for *S. palustre* and *S. squarrosum.* Besides sucrose, glucose and fructose could be detected right at the beginning of the cultivation (Figure 3-5, C-E). The disaccharide sucrose partially hydrolysed to its monosaccharides glucose and fructose due to high temperature during autoclaving as reported before (Ball, 1953).

The initial sugar concentration was higher than expected for *S. palustre* and *S. squarrosum*. During the hydrolysis of sucrose a water molecule is added, which results in a 5 % higher molecular weight of the monosaccharides (glucose/fructose 180.16 g mol^−1^) compared to the disaccharide (sucrose 342.30 g mol^−1^). Due to the low standard deviation of the sugar measurements, another reason for the concentration difference could be the preculture medium still containing not completely utilized sugars, which would increase the available amount at the beginning of the bioreactor cultivation. The growth of the preculture seems to be slower compared to the bioreactor culture due to different aeration without CO_2_ supplementation, different light sources resulting in different light spectra and lower light availibilty. On the other hand, it is reported that light intensity had no influence on the growth of several moss species in organic nutrient medium (Fries, 1945) and on the growth of *S. nemoreum* in the presence of exogenous sugars, but the light quality and quantity used was probably not the same (Simola, 1969).

For the initial high sucrose content it is also possible that the sugar assay is not absolutely specific for sucrose, as β-fructosidase also hydrolyses low molecular weight fructans (Megazyme Booklet), which could increase the amount of the measured sucrose. Sucrose and fructan are the major soluble carbohydrates of *Sphagnum* (Maass and Craigie, 1964; Marschall and Laufer, 2002). The secretion of fructan by peat mosses has not been reported yet, but they contribute to the total dissolved organic carbon of peat leachate (Fenner et al., 2004).

Further analysis of sugar concentrations in our cultivation media showed that during 10 to 13 days of cultivation of *S. palustre* and *S. fuscum*, nearly all sucrose was depleted from the medium, whereas the concentrations of glucose and fructose increased during that time. The pH courses (Figure 3-5, B) indicate that hydrolysis did not occur due to acidification of the medium, because the pH minimum was reached at day 3 of cultivation, while the sucrose was depleted approximately on day 10, dependent on the species. We suppose that most of the sucrose is hydrolysed by enzymes secreted by the mosses. In accordance with this are reports on sucrose cleavage by cell wall acid-type invertases in *S. nemoreum* (Simola, 1969), in *S. recurvum* (Marschall and Laufer, 2002) and in *S. compactum* (Graham et al., 2010). Comparing the sucrose concentration during the first three days of cultivation of *S. squarrosum* with *S. fuscum* suggests that *S. squarrosum* is capable of cleaving less sucrose compared to *S. fuscum* under the same pH changes. This could be the result of different enzyme activities of invertases among the species. Future analyses may reveal whether the moss species differ in the enzyme activities of acid-invertases or in enzyme amounts in the cell wall.

After the absence of detectable sucrose, glucose and fructose concentrations were decreasing until both sugars were consumed. We observed that the total consumption of fructose took place two days later compared to glucose in *S. palustre*’s cultivation. At the end of *S. fuscum*’s cultivation, glucose was nearly depleted and up to 2 g L^−1^ of fructose remained. The ablity to take up exogenuous sugars is well known for *Sphagnum* mosses (Simola, 1969, Graham et al., 2010). Glucose and fructose can be absorbed via monossacharide transporters and utilized for growth and energy (Simola 1969). Graham et al. (2010) showed that glucose is preferentially taken up by *Sphagnum* mosses, which correlates with the findings in our study, because glucose was absorbed more quickly than fructose.

Our results also show that *S. squarrosum* used organic carbon less effectively than both other species, because some sucrose was still left in the medium at the end of the cultivation (Figure 5, C). The same amount of added sucrose generated 34 % less biomass of *S. squarrosum* than *S. palustre*. Apart from the possible lower enzyme activity of acid-invertase in this species, the overreaching of pH 5.5 may also affect the hydrolysis. Due to the fact that the invertase enzymes have their optimum activity at pH values between 4.0 and 5.5 (Chibbar et al., 2016), the sucrose hydrolysis could be partially slowed down with increasing pH. *S. squarrosum* is able to tolerate base-rich water (Daniels and Eddy, 1990) and is less sensitive to higher pH than *S. palustre* is (Clymo, 1973; Harpenslager et al., 2015).

As sucrose is not completely utilized, but has a positive effect on the growth of *S. squarrosum*, the presence of sucrose might have an indirect effect on the growth. Sugars are not only important in plant energy metabolism, they are also important signal molecules that interact with several hormones and serve as morphogens (Rolland et al., 2002). Furthermore, the vacuolar osmotic potential is altered by polymerization or breakdown of fructan and this may alter turgor pressure (Marschall, 2010), which may improve the nutrient uptake. Such questions may be resolved in future studies using transcriptome profiling, as has been demonstrated for seed plants (e.g., Wang et al., 2018).

## 4. Conclusion

The major aim of this study was to optimize the culture media for increased biomass productivity of three *Sphagnum* species using fractional factorial and central composite design. The media optimization in flask cultures allowed us to lower nutrient concentrations while achieving better productivity. This made upscaling in bioreactors feasible. The lower demand on nutrients in the optimized culture media makes the production process economically favorable to implement the next step towards large scale *Sphagnum* biomass production. Furthermore, other process parameters like light intensity and CO_2_ supply can be optimized in future by this tool.

## 5. Acknowledgements

This work was funded by the Federal Ministry of Food and Agriculture (BMEL) (MOOSzucht, No. 22007216 to C.P. and R.R.). Additional support came from the German Research Foundation (DFG) under Germany’s Excellence Strategy (CIBSS – EXC-2189 – Project ID 390939984 and *liv*MatS – EXC-2193 to R.R.). We gratefully acknowledge Anja Kuberski for technical assistance, Olga Gorte and Christin Kubisch for performing HPLC measurements and Anne Katrin Prowse for proofreading of the manuscript.

## 6. Author’s contribution

Melanie Heck: Investigation, Methodology, Validation, Formal analysis, Writing – Original draft preparation. Ingrida Melková: Investigation, Methodology, Writing – Review & Editing. Clemens Posten: Supervision, Writing – Review & Editing, Funding Acquisition. Eva Decker: Conceptualization, Supervision, Writing – Review & Editing. Ralf Reski: Supervision, Writing – Review & Editing, Funding Acquisition.

## 8. Supplement

**Table S1:** Experimental design and results of the two-level factorial design used as preliminary screening for the optimized media composition of *S. fuscum* sorted by the standard (Std) order of the design matrix including two runs of the factorial design and four center points. The number of runs was conducted in random order. The factors A (Sucrose in g L^−1^), B (NH4NO3 in mM), C (KH2PO4 in mM), D (KCl in mM), E (MgSO4 in mM), F (Ca(NO3)2 in mM), G (ME in %) and H (FeSO4 in µM) were set at the three levels: reduced and elevated concentration and in between. The biomass after four weeks of cultivation was measured in mg dry weight. The chemical formulas of hydrated salts are expressed without water molecules.

| Std | Run | A | B | C | D | E | F | G | H | Response | Average Response |
| --- | --- | --- | --- | --- | --- | --- | --- | --- | --- | --- | --- |
|  |  | Sucrose<br>g/l | NH<sub>4</sub>NO<sub>3</sub><br>mM | KH<sub>2</sub>PO<sub>4</sub><br>mM | KCl<br>mM | MgSO<sub>4</sub><br>mM | Ca(NO<sub>3</sub>)<sub>2</sub><br>mM | ME<br>% | FeSO<sub>4</sub><br>μM | Biomass<br>mg DW | Biomass<br>mg DW |
| 1 | 32 | 3 | 0.63 | 0.92 | 1.68 | 0.51 | 2.12 | 50 | 22.5 | 40.79 | 39.42 ± 1.38 |
| 2 | 31 | 3 | 0.63 | 0.92 | 1.68 | 0.51 | 2.12 | 50 | 22.5 | 38.04 |  |
| 3 | 27 | 20 | 0.63 | 0.92 | 1.68 | 1.01 | 2.12 | 100 | 45 | 44.52 | 52.90 ± 8.38 |
| 4 | 26 | 20 | 0.63 | 0.92 | 1.68 | 1.01 | 2.12 | 100 | 45 | 61.28 |  |
| 5 | 13 | 3 | 1.25 | 0.92 | 1.68 | 1.01 | 4.24 | 50 | 45 | 33.35 | 39.95 ± 6.59 |
| 6 | 17 | 3 | 1.25 | 0.92 | 1.68 | 1.01 | 4.24 | 50 | 45 | 46.54 |  |
| 7 | 6 | 20 | 1.25 | 0.92 | 1.68 | 0.51 | 4.24 | 100 | 22.5 | 57.32 | 61.44 ± 4.12 |
| 8 | 20 | 20 | 1.25 | 0.92 | 1.68 | 0.51 | 4.24 | 100 | 22.5 | 65.55 |  |
| 9 | 14 | 3 | 0.63 | 1.84 | 1.68 | 1.01 | 4.24 | 100 | 22.5 | 45.45 | 43.42 ± 2.04 |
| 10 | 29 | 3 | 0.63 | 1.84 | 1.68 | 1.01 | 4.24 | 100 | 22.5 | 41.38 |  |
| 11 | 11 | 20 | 0.63 | 1.84 | 1.68 | 0.51 | 4.24 | 50 | 45 | 53.41 | 56.79 ± 3.38 |
| 12 | 16 | 20 | 0.63 | 1.84 | 1.68 | 0.51 | 4.24 | 50 | 45 | 60.17 |  |
| 13 | 21 | 3 | 1.25 | 1.84 | 1.68 | 0.51 | 2.12 | 100 | 45 | 50.06 | 53.97 ± 3.91 |
| 14 | 34 | 3 | 1.25 | 1.84 | 1.68 | 0.51 | 2.12 | 100 | 45 | 57.88 |  |
| 15 | 24 | 20 | 1.25 | 1.84 | 1.68 | 1.01 | 2.12 | 50 | 22.5 | 62.02 | 54.48 ± 7.55 |
| 16 | 3 | 20 | 1.25 | 1.84 | 1.68 | 1.01 | 2.12 | 50 | 22.5 | 46.93 |  |
| 17 | 19 | 3 | 0.63 | 0.92 | 3.35 | 0.51 | 4.24 | 100 | 45 | 43.82 | 42.22 ± 1.60 |
| 18 | 23 | 3 | 0.63 | 0.92 | 3.35 | 0.51 | 4.24 | 100 | 45 | 40.62 |  |
| 19 | 33 | 20 | 0.63 | 0.92 | 3.35 | 1.01 | 4.24 | 50 | 22.5 | 62.6 | 54.53 ± 8.08 |
| 20 | 5 | 20 | 0.63 | 0.92 | 3.35 | 1.01 | 4.24 | 50 | 22.5 | 46.45 |  |
| 21 | 30 | 3 | 1.25 | 0.92 | 3.35 | 1.01 | 2.12 | 100 | 22.5 | 39.81 | 43.62 ± 3.81 |
| 22 | 10 | 3 | 1.25 | 0.92 | 3.35 | 1.01 | 2.12 | 100 | 22.5 | 47.42 |  |
| 23 | 15 | 20 | 1.25 | 0.92 | 3.35 | 0.51 | 2.12 | 50 | 45 | 47.93 | 50.46 ± 2.53 |
| 24 | 8 | 20 | 1.25 | 0.92 | 3.35 | 0.51 | 2.12 | 50 | 45 | 52.98 |  |
| 25 | 18 | 3 | 0.63 | 1.84 | 3.35 | 1.01 | 2.12 | 50 | 45 | 45.57 | 42.91 ± 2.67 |
| 26 | 9 | 3 | 0.63 | 1.84 | 3.35 | 1.01 | 2.12 | 50 | 45 | 40.24 |  |
| 27 | 28 | 20 | 0.63 | 1.84 | 3.35 | 0.51 | 2.12 | 100 | 22.5 | 53.03 | 53.58 ± 0.54 |
| 28 | 36 | 20 | 0.63 | 1.84 | 3.35 | 0.51 | 2.12 | 100 | 22.5 | 54.12 |  |
| 29 | 1 | 3 | 1.25 | 1.84 | 3.35 | 0.51 | 4.24 | 50 | 22.5 | 44.18 | 45.15 ± 0.97 |
| 30 | 25 | 3 | 1.25 | 1.84 | 3.35 | 0.51 | 4.24 | 50 | 22.5 | 46.11 |  |
| 31 | 22 | 20 | 1.25 | 1.84 | 3.35 | 1.01 | 4.24 | 100 | 45 | 61.02 | 62.62 ± 1.60 |
| 32 | 35 | 20 | 1.25 | 1.84 | 3.35 | 1.01 | 4.24 | 100 | 45 | 64.22 |  |
| 33 | 7 | 11.5 | 0.94 | 1.38 | 2.51 | 0.76 | 3.18 | 75 | 33.75 | 49.89 | 55.10 ± 4.25 |
| 34 | 12 | 11.5 | 0.94 | 1.38 | 2.51 | 0.76 | 3.18 | 75 | 33.75 | 54.01 |  |
| 35 | 2 | 11.5 | 0.94 | 1.38 | 2.51 | 0.76 | 3.18 | 75 | 33.75 | 61.71 |  |
| 36 | 4 | 11.5 | 0.94 | 1.38 | 2.51 | 0.76 | 3.18 | 75 | 33.75 | 54.79 |  |

**Table S2:**
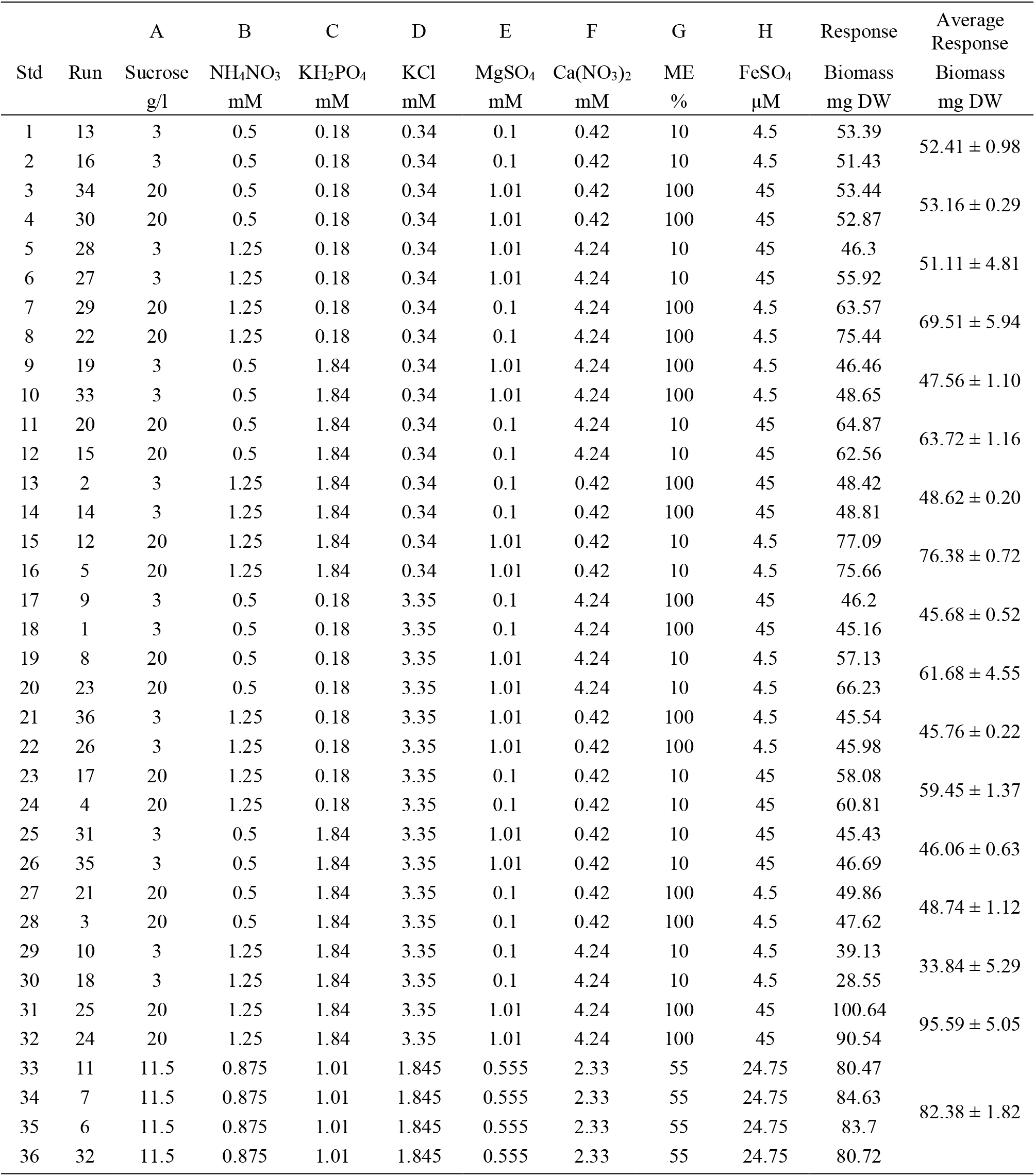
Experimental design and results of the two-level factorial design used as preliminary screening for the optimized media composition of *S. palustre* sorted by the standard (Std) order of the design matrix including two runs of the factorial design and four center points. The number of runs was conducted in random order. The factors A (Sucrose in g L^−1^), B (NH4NO3 in mM), C (KH2PO4 in mM), D (KCl in mM), E (MgSO4 in mM), F (Ca(NO3)2 in mM), G (ME in %) and H (FeSO4 in µM) were set at the three levels: reduced and elevated concentration and in between. The biomass after four weeks of cultivation was measured in mg dry weight. The chemical formulas of hydrated salts are expressed without water molecules.

**Table S3:** Experimental design and results of the two-level factorial design used as preliminary screening for the optimized media composition of *S. squarrosum* sorted by the standard (Std) order of the design matrix including two runs of the factorial design and four center points. The number of runs was conducted in random order. The factors A (Sucrose in g L^−1^), B (NH4NO3 in mM), C (KH2PO4 in mM), D (KCl in mM), E (MgSO4 in mM), F (Ca(NO3)2 in mM), G (ME in %) and H (FeSO4 in µM) were set at the three levels: reduced and elevated concentration and in between. The biomass after four weeks of cultivation was measured in mg dry weight. The chemical formulas of hydrated salts are expressed without water molecules.

| Std | Run | A | B | C | D | E | F | G | H | Response | Average Response |
| --- | --- | --- | --- | --- | --- | --- | --- | --- | --- | --- | --- |
|  |  | Sucrose<br>g/l | NH<sub>4</sub>NO<sub>3</sub><br>mM | KH<sub>2</sub>PO<sub>4</sub><br>mM | KCl<br>mM | MgSO<sub>4</sub><br>mM | Ca(NO<sub>3</sub>)<sub>2</sub><br>mM | ME<br>% | FeSO<sub>4</sub><br>μM | Biomass<br>mg DW | Biomass<br>mg DW |
| 1 | 32 | 3 | 0.63 | 0.92 | 1.68 | 0.51 | 2.12 | 50 | 22.5 | 18.55 | 24.89 ± 6.34 |
| 2 | 31 | 3 | 0.63 | 0.92 | 1.68 | 0.51 | 2.12 | 50 | 22.5 | 31.22 |  |
| 3 | 27 | 20 | 0.63 | 0.92 | 1.68 | 1.01 | 2.12 | 100 | 45 | 17.7 | 20.13 ± 2.43 |
| 4 | 26 | 20 | 0.63 | 0.92 | 1.68 | 1.01 | 2.12 | 100 | 45 | 22.55 |  |
| 5 | 13 | 3 | 1.25 | 0.92 | 1.68 | 1.01 | 4.24 | 50 | 45 | 24.85 | 27.92 ± 3.07 |
| 6 | 17 | 3 | 1.25 | 0.92 | 1.68 | 1.01 | 4.24 | 50 | 45 | 30.98 |  |
| 7 | 6 | 20 | 1.25 | 0.92 | 1.68 | 0.51 | 4.24 | 100 | 22.5 | 33.69 | 29.99 ± 3.71 |
| 8 | 20 | 20 | 1.25 | 0.92 | 1.68 | 0.51 | 4.24 | 100 | 22.5 | 26.28 |  |
| 9 | 14 | 3 | 0.63 | 1.84 | 1.68 | 1.01 | 4.24 | 100 | 22.5 | 20.25 | 23.75 ± 3.50 |
| 10 | 29 | 3 | 0.63 | 1.84 | 1.68 | 1.01 | 4.24 | 100 | 22.5 | 27.24 |  |
| 11 | 11 | 20 | 0.63 | 1.84 | 1.68 | 0.51 | 4.24 | 50 | 45 | 45.55 | 40.43 ± 5.13 |
| 12 | 16 | 20 | 0.63 | 1.84 | 1.68 | 0.51 | 4.24 | 50 | 45 | 35.3 |  |
| 13 | 21 | 3 | 1.25 | 1.84 | 1.68 | 0.51 | 2.12 | 100 | 45 | 14.09 | 15.01 ± 0.92 |
| 14 | 34 | 3 | 1.25 | 1.84 | 1.68 | 0.51 | 2.12 | 100 | 45 | 15.93 |  |
| 15 | 24 | 20 | 1.25 | 1.84 | 1.68 | 1.01 | 2.12 | 50 | 22.5 | 25.67 | 28.08 ± 2.41 |
| 16 | 3 | 20 | 1.25 | 1.84 | 1.68 | 1.01 | 2.12 | 50 | 22.5 | 30.48 |  |
| 17 | 19 | 3 | 0.63 | 0.92 | 3.35 | 0.51 | 4.24 | 100 | 45 | 18.92 | 19.97 ± 1.05 |
| 18 | 23 | 3 | 0.63 | 0.92 | 3.35 | 0.51 | 4.24 | 100 | 45 | 21.02 |  |
| 19 | 33 | 20 | 0.63 | 0.92 | 3.35 | 1.01 | 4.24 | 50 | 22.5 | 39.99 | 34.96 ± 5.04 |
| 20 | 5 | 20 | 0.63 | 0.92 | 3.35 | 1.01 | 4.24 | 50 | 22.5 | 29.92 |  |
| 21 | 30 | 3 | 1.25 | 0.92 | 3.35 | 1.01 | 2.12 | 100 | 22.5 | 15.05 | 15.45 ± 0.40 |
| 22 | 10 | 3 | 1.25 | 0.92 | 3.35 | 1.01 | 2.12 | 100 | 22.5 | 15.85 |  |
| 23 | 15 | 20 | 1.25 | 0.92 | 3.35 | 0.51 | 2.12 | 50 | 45 | 19.42 | 14.77 ± 4.66 |
| 24 | 8 | 20 | 1.25 | 0.92 | 3.35 | 0.51 | 2.12 | 50 | 45 | 10.11 |  |
| 25 | 18 | 3 | 0.63 | 1.84 | 3.35 | 1.01 | 2.12 | 50 | 45 | 36.42 | 34.66 ± 1.77 |
| 26 | 9 | 3 | 0.63 | 1.84 | 3.35 | 1.01 | 2.12 | 50 | 45 | 32.89 |  |
| 27 | 28 | 20 | 0.63 | 1.84 | 3.35 | 0.51 | 2.12 | 100 | 22.5 | 24.02 | 25.80 ± 1.78 |
| 28 | 36 | 20 | 0.63 | 1.84 | 3.35 | 0.51 | 2.12 | 100 | 22.5 | 27.57 |  |
| 29 | 1 | 3 | 1.25 | 1.84 | 3.35 | 0.51 | 4.24 | 50 | 22.5 | 25.41 | 22.72 ± 2.69 |
| 30 | 25 | 3 | 1.25 | 1.84 | 3.35 | 0.51 | 4.24 | 50 | 22.5 | 20.03 |  |
| 31 | 22 | 20 | 1.25 | 1.84 | 3.35 | 1.01 | 4.24 | 100 | 45 | 24.44 | 21.23 ± 3.21 |
| 32 | 35 | 20 | 1.25 | 1.84 | 3.35 | 1.01 | 4.24 | 100 | 45 | 18.02 |  |
| 33 | 7 | 11.5 | 0.94 | 1.38 | 2.51 | 0.76 | 3.18 | 75 | 33.75 | 29.74 | 36.45 ± 5.73 |
| 34 | 12 | 11.5 | 0.94 | 1.38 | 2.51 | 0.76 | 3.18 | 75 | 33.75 | 33.26 |  |
| 35 | 2 | 11.5 | 0.94 | 1.38 | 2.51 | 0.76 | 3.18 | 75 | 33.75 | 45.09 |  |
| 36 | 4 | 11.5 | 0.94 | 1.38 | 2.51 | 0.76 | 3.18 | 75 | 33.75 | 37.72 |  |

**Table S4:** Experimental design and results of the central composite design for optimization of the media composition of *S. fuscum* sorted by the standard (Std) order of the design matrix including the runs for factorial points, ten axial points and eight center points. The number of runs was conducted in random order. The factors A (Sucrose in g L^−1^), B (NH4NO3 in mM), C (KH2PO4 in mM), D (Ca(NO3)2 in mM) and E (ME in %) were set at five levels: axial points, reduced and elevated concentration and in between. The biomass after four weeks of cultivation was measured in mg dry weight. The chemical formulas of hydrated salts are expressed without water molecules.

| Std | Run | A<br>Sucrose<br>g/L | B<br>NH<sub>4</sub>NO<sub>3</sub><br>mM | C<br>KH<sub>2</sub>PO<sub>4</sub><br>mM | D<br>Ca(NO<sub>3</sub>)<sub>2</sub><br>mM | E<br>ME<br>% | Response<br>Biomass<br>mg DW | Predicted<br>Biomass<br>mg DW |
| --- | --- | --- | --- | --- | --- | --- | --- | --- |
| 1 | 32 | 5 | 0.63 | 0.92 | 2.12 | 50 | 403.3 | 334.7 |
| 2 | 29 | 20 | 0.63 | 0.92 | 2.12 | 50 | 409.3 | 430.3 |
| 3 | 40 | 5 | 1.25 | 0.92 | 2.12 | 50 | 362.6 | 319.5 |
| 4 | 25 | 20 | 1.25 | 0.92 | 2.12 | 50 | 451.4 | 408.2 |
| 5 | 49 | 5 | 0.63 | 1.84 | 2.12 | 50 | 292.2 | 315.3 |
| 6 | 27 | 20 | 0.63 | 1.84 | 2.12 | 50 | 404.9 | 402.2 |
| 7 | 22 | 5 | 1.25 | 1.84 | 2.12 | 50 | 257 | 301.4 |
| 8 | 5 | 20 | 1.25 | 1.84 | 2.12 | 50 | 356.6 | 382.2 |
| 9 | 20 | 5 | 0.63 | 0.92 | 4.24 | 50 | 288.4 | 306.5 |
| 10 | 26 | 20 | 0.63 | 0.92 | 4.24 | 50 | 365.9 | 389.5 |
| 11 | 44 | 5 | 1.25 | 0.92 | 4.24 | 50 | 351.8 | 293.2 |
| 12 | 41 | 20 | 1.25 | 0.92 | 4.24 | 50 | 322 | 370.4 |
| 13 | 19 | 5 | 0.63 | 1.84 | 4.24 | 50 | 262.5 | 289.5 |
| 14 | 6 | 20 | 0.63 | 1.84 | 4.24 | 50 | 416.7 | 365.2 |
| 15 | 39 | 5 | 1.25 | 1.84 | 4.24 | 50 | 274.4 | 277.2 |
| 16 | 21 | 20 | 1.25 | 1.84 | 4.24 | 50 | 311.9 | 347.9 |
| 17 | 2 | 5 | 0.63 | 0.92 | 2.12 | 100 | 357.9 | 334.7 |
| 18 | 4 | 20 | 0.63 | 0.92 | 2.12 | 100 | 345.6 | 430.3 |
| 19 | 1 | 5 | 1.25 | 0.92 | 2.12 | 100 | 330.4 | 319.5 |
| 20 | 18 | 20 | 1.25 | 0.92 | 2.12 | 100 | 361.2 | 408.2 |
| 21 | 35 | 5 | 0.63 | 1.84 | 2.12 | 100 | 347.1 | 315.3 |
| 22 | 15 | 20 | 0.63 | 1.84 | 2.12 | 100 | 461.7 | 402.2 |
| 23 | 50 | 5 | 1.25 | 1.84 | 2.12 | 100 | 310.9 | 301.4 |
| 24 | 47 | 20 | 1.25 | 1.84 | 2.12 | 100 | 379.6 | 382.2 |
| 25 | 45 | 5 | 0.63 | 0.92 | 4.24 | 100 | 262.3 | 306.5 |
| 26 | 31 | 20 | 0.63 | 0.92 | 4.24 | 100 | 417.2 | 389.5 |
| 27 | 34 | 5 | 1.25 | 0.92 | 4.24 | 100 | 362.3 | 293.2 |
| 28 | 42 | 20 | 1.25 | 0.92 | 4.24 | 100 | 379.7 | 370.4 |
| 29 | 8 | 5 | 0.63 | 1.84 | 4.24 | 100 | 284.4 | 289.5 |
| 30 | 9 | 20 | 0.63 | 1.84 | 4.24 | 100 | 403.3 | 365.2 |
| 31 | 43 | 5 | 1.25 | 1.84 | 4.24 | 100 | 288.1 | 277.2 |
| 32 | 10 | 20 | 1.25 | 1.84 | 4.24 | 100 | 338.5 | 347.9 |
| 33 | 3 | 1.28 | 0.94 | 1.38 | 3.18 | 75 | 209.8 | 281.7 |
| 34 | 23 | 23.72 | 0.94 | 1.38 | 3.18 | 75 | 415.8 | 401.1 |
| 35 | 30 | 12.5 | 0.47 | 1.38 | 3.18 | 75 | 352.2 | 402.8 |
| 36 | 46 | 12.5 | 1.40 | 1.38 | 3.18 | 75 | 308.6 | 373.4 |
| 37 | 28 | 12.5 | 0.94 | 0.69 | 3.18 | 75 | 316.7 | 411.6 |
| 38 | 12 | 12.5 | 0.94 | 2.07 | 3.18 | 75 | 349.3 | 373.1 |
| 39 | 7 | 12.5 | 0.94 | 1.38 | 1.59 | 75 | 405.7 | 516.8 |
| 40 | 36 | 12.5 | 0.94 | 1.38 | 4.77 | 75 | 359.8 | 439.9 |
| 41 | 17 | 12.5 | 0.94 | 1.38 | 3.18 | 37.6 | 350 | 476.0 |
| 42 | 37 | 12.5 | 0.94 | 1.38 | 3.18 | 123.8 | 367.3 | 476.0 |
| 43 | 11 | 12.5 | 0.94 | 1.38 | 3.18 | 75 | 542.7 | 476.0 |
| 44 | 14 | 12.5 | 0.94 | 1.38 | 3.18 | 75 | 569.1 | 476.0 |
| 45 | 38 | 12.5 | 0.94 | 1.38 | 3.18 | 75 | 555.2 | 476.0 |
| 46 | 16 | 12.5 | 0.94 | 1.38 | 3.18 | 75 | 560.6 | 476.0 |
| 47 | 48 | 12.5 | 0.94 | 1.38 | 3.18 | 75 | 595.8 | 475.0 |
| 48 | 24 | 12.5 | 0.94 | 1.38 | 3.18 | 75 | 557.8 | 476.0 |
| 49 | 33 | 12.5 | 0.94 | 1.38 | 3.18 | 75 | 455.8 | 476.0 |
| 50 | 13 | 12.5 | 0.94 | 1.38 | 3.18 | 75 | 544.8 | 476.0 |

**Table S5:**
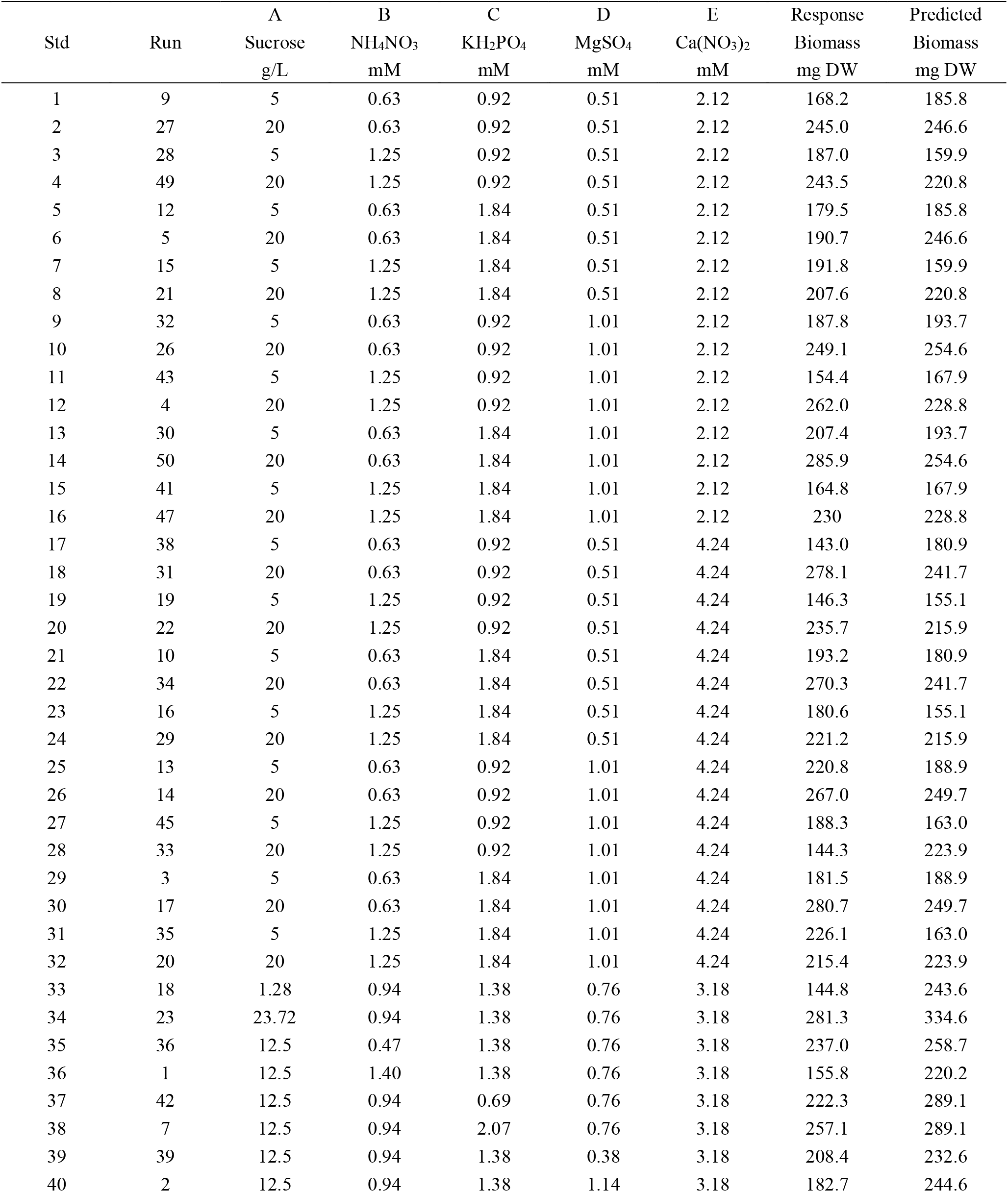
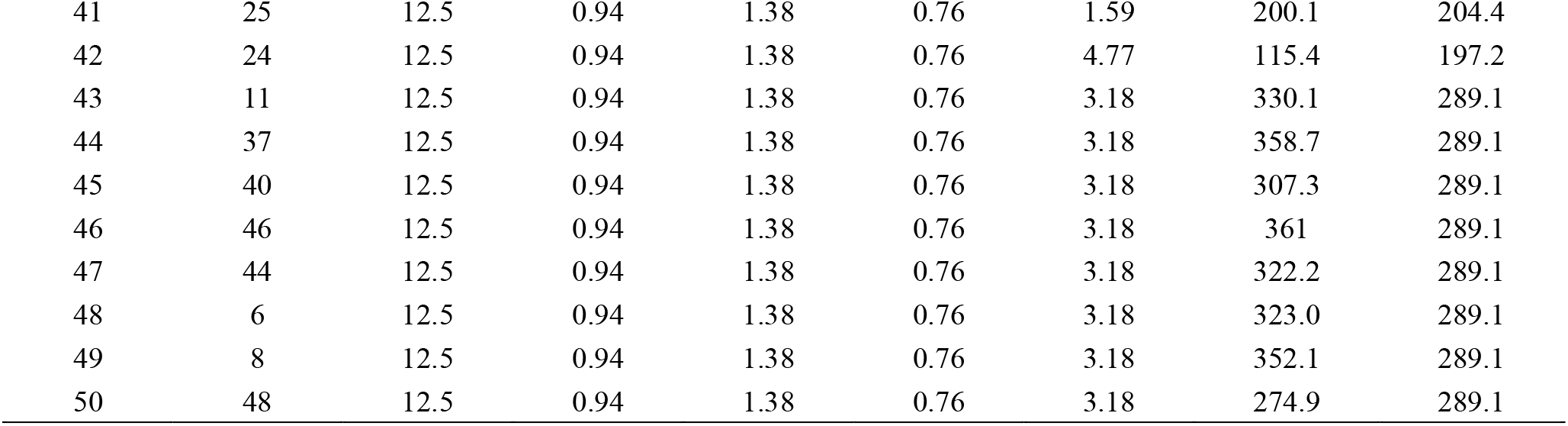
Experimental design and results of the central composite design for optimization of the media composition of *S. palustre* sorted by the standard (Std) order of the design matrix including the runs for factorial points, ten axial points and eight center points. The number of runs was conducted in random order. The factors A (Sucrose in g L^−1^), B (NH4NO3 in mM), C (KH2PO4 in mM), D (MgSO4 in mM) and (Ca(NO3)2 in mM) were set at five levels: axial points, reduced and elevated concentration and in between. The biomass after four weeks of cultivation was measured in mg dry weight. The chemical formulas of hydrated salts are expressed without water molecules.

**Table S6:**
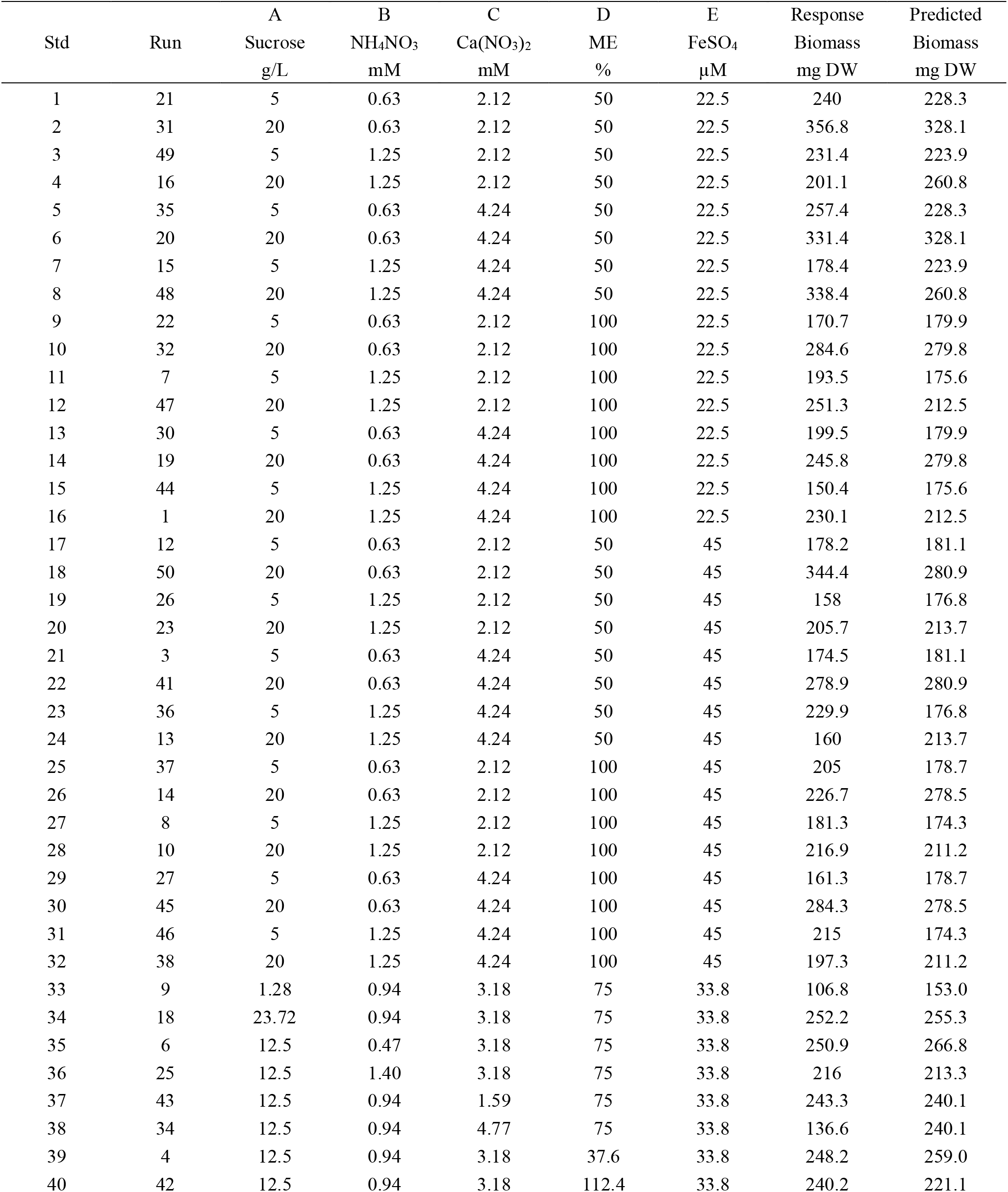
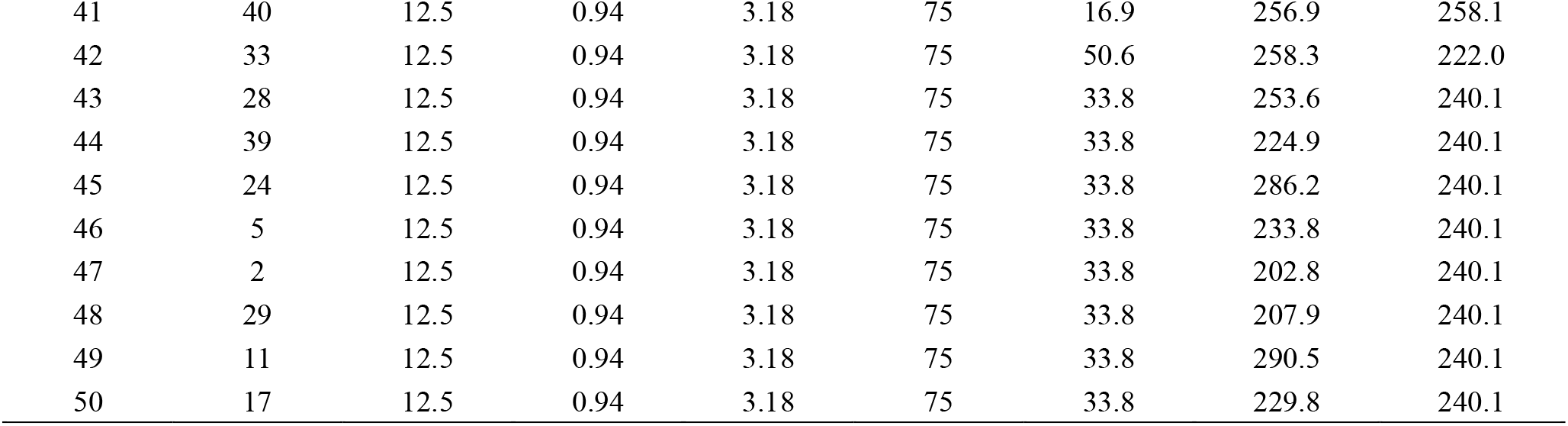
Experimental design and results of the central composite design for optimization of the media composition of *S. squarrosum* sorted by the standard (Std) order of the design matrix including the runs for factorial points, ten axial points and eight center points. The number of runs was conducted in random order. The factors A (Sucrose in g L^−1^), B (NH4NO3 in mM), C (Ca(NO3)2 in mM), D (ME in %) and E (FeSO4 in µM) were set at five levels: axial points, reduced and elevated concentration and in between. The biomass after five weeks of cultivation was measured in mg dry weight. The chemical formulas of hydrated salts are expressed without water molecules.

| Std | Run | A<br>Sucrose<br>g/L | B<br>NH<sub>4</sub>NO<sub>3</sub><br>mM | C<br>Ca(NO<sub>3</sub>)<sub>2</sub><br>mM | D<br>ME<br>% | E<br>FeSO<sub>4</sub><br>μM | Response<br>Biomass<br>mg DW | Predicted<br>Biomass<br>mg DW |
| --- | --- | --- | --- | --- | --- | --- | --- | --- |
| 1 | 21 | 5 | 0.63 | 2.12 | 50 | 22.5 | 240 | 228.3 |
| 2 | 31 | 20 | 0.63 | 2.12 | 50 | 22.5 | 356.8 | 328.1 |
| 3 | 49 | 5 | 1.25 | 2.12 | 50 | 22.5 | 231.4 | 223.9 |
| 4 | 16 | 20 | 1.25 | 2.12 | 50 | 22.5 | 201.1 | 260.8 |
| 5 | 35 | 5 | 0.63 | 4.24 | 50 | 22.5 | 257.4 | 228.3 |
| 6 | 20 | 20 | 0.63 | 4.24 | 50 | 22.5 | 331.4 | 328.1 |
| 7 | 15 | 5 | 1.25 | 4.24 | 50 | 22.5 | 178.4 | 223.9 |
| 8 | 48 | 20 | 1.25 | 4.24 | 50 | 22.5 | 338.4 | 260.8 |
| 9 | 22 | 5 | 0.63 | 2.12 | 100 | 22.5 | 170.7 | 179.9 |
| 10 | 32 | 20 | 0.63 | 2.12 | 100 | 22.5 | 284.6 | 279.8 |
| 11 | 7 | 5 | 1.25 | 2.12 | 100 | 22.5 | 193.5 | 175.6 |
| 12 | 47 | 20 | 1.25 | 2.12 | 100 | 22.5 | 251.3 | 212.5 |
| 13 | 30 | 5 | 0.63 | 4.24 | 100 | 22.5 | 199.5 | 179.9 |
| 14 | 19 | 20 | 0.63 | 4.24 | 100 | 22.5 | 245.8 | 279.8 |
| 15 | 44 | 5 | 1.25 | 4.24 | 100 | 22.5 | 150.4 | 175.6 |
| 16 | 1 | 20 | 1.25 | 4.24 | 100 | 22.5 | 230.1 | 212.5 |
| 17 | 12 | 5 | 0.63 | 2.12 | 50 | 45 | 178.2 | 181.1 |
| 18 | 50 | 20 | 0.63 | 2.12 | 50 | 45 | 344.4 | 280.9 |
| 19 | 26 | 5 | 1.25 | 2.12 | 50 | 45 | 158 | 176.8 |
| 20 | 23 | 20 | 1.25 | 2.12 | 50 | 45 | 205.7 | 213.7 |
| 21 | 3 | 5 | 0.63 | 4.24 | 50 | 45 | 174.5 | 181.1 |
| 22 | 41 | 20 | 0.63 | 4.24 | 50 | 45 | 278.9 | 280.9 |
| 23 | 36 | 5 | 1.25 | 4.24 | 50 | 45 | 229.9 | 176.8 |
| 24 | 13 | 20 | 1.25 | 4.24 | 50 | 45 | 160 | 213.7 |
| 25 | 37 | 5 | 0.63 | 2.12 | 100 | 45 | 205 | 178.7 |
| 26 | 14 | 20 | 0.63 | 2.12 | 100 | 45 | 226.7 | 278.5 |
| 27 | 8 | 5 | 1.25 | 2.12 | 100 | 45 | 181.3 | 174.3 |
| 28 | 10 | 20 | 1.25 | 2.12 | 100 | 45 | 216.9 | 211.2 |
| 29 | 27 | 5 | 0.63 | 4.24 | 100 | 45 | 161.3 | 178.7 |
| 30 | 45 | 20 | 0.63 | 4.24 | 100 | 45 | 284.3 | 278.5 |
| 31 | 46 | 5 | 1.25 | 4.24 | 100 | 45 | 215 | 174.3 |
| 32 | 38 | 20 | 1.25 | 4.24 | 100 | 45 | 197.3 | 211.2 |
| 33 | 9 | 1.28 | 0.94 | 3.18 | 75 | 33.8 | 106.8 | 153.0 |
| 34 | 18 | 23.72 | 0.94 | 3.18 | 75 | 33.8 | 252.2 | 255.3 |
| 35 | 6 | 12.5 | 0.47 | 3.18 | 75 | 33.8 | 250.9 | 266.8 |
| 36 | 25 | 12.5 | 1.40 | 3.18 | 75 | 33.8 | 216 | 213.3 |
| 37 | 43 | 12.5 | 0.94 | 1.59 | 75 | 33.8 | 243.3 | 240.1 |
| 38 | 34 | 12.5 | 0.94 | 4.77 | 75 | 33.8 | 136.6 | 240.1 |
| 39 | 4 | 12.5 | 0.94 | 3.18 | 37.6 | 33.8 | 248.2 | 259.0 |
| 40 | 42 | 12.5 | 0.94 | 3.18 | 112.4 | 33.8 | 240.2 | 221.1 |
| 41 | 40 | 12.5 | 0.94 | 3.18 | 75 | 16.9 | 256.9 | 258.1 |
| 42 | 33 | 12.5 | 0.94 | 3.18 | 75 | 50.6 | 258.3 | 222.0 |
| 43 | 28 | 12.5 | 0.94 | 3.18 | 75 | 33.8 | 253.6 | 240.1 |
| 44 | 39 | 12.5 | 0.94 | 3.18 | 75 | 33.8 | 224.9 | 240.1 |
| 45 | 24 | 12.5 | 0.94 | 3.18 | 75 | 33.8 | 286.2 | 240.1 |
| 46 | 5 | 12.5 | 0.94 | 3.18 | 75 | 33.8 | 233.8 | 240.1 |
| 47 | 2 | 12.5 | 0.94 | 3.18 | 75 | 33.8 | 202.8 | 240.1 |
| 48 | 29 | 12.5 | 0.94 | 3.18 | 75 | 33.8 | 207.9 | 240.1 |
| 49 | 11 | 12.5 | 0.94 | 3.18 | 75 | 33.8 | 290.5 | 240.1 |
| 50 | 17 | 12.5 | 0.94 | 3.18 | 75 | 33.8 | 229.8 | 240.1 |

## Notes

### Competing Interest Statement

The authors have declared no competing interest.

